# Development of prefrontal corticostriatal connectivity in mice

**DOI:** 10.1101/2023.03.14.532475

**Authors:** Roxana E. Mesías, Yosif Zaki, Christopher A. Guevara, Lauren G. Friedman, Ayan Hussein, Karen Therrien, Alexandra R. Magee, Nikolaos Tzavaras, Pamela Del Valle, Mark G. Baxter, George W. Huntley, Deanna L. Benson

**Affiliations:** Nash Family Department of Neuroscience, Friedman Brain Institute, Graduate School of Biomedical Sciences, Icahn School of Medicine at Mount Sinai, New York, NY 10029

**Keywords:** prefrontal cortex, dorsomedial striatum, cadherin-8, synaptogenesis, electrophysiology, instrumental conditioning, optogenetics

## Abstract

Rational decision making is grounded in learning to associate actions with outcomes, a process that depends on projections from prefrontal cortex to dorsomedial striatum. Symptoms associated with a variety of human pathological conditions ranging from schizophrenia and autism to Huntington’s and Parkinson’s disease point toward functional deficits in this projection, but its development is not well understood, making it difficult to investigate how perturbations in development of this circuitry could contribute to pathophysiology. We applied a novel strategy based on Hotspot Analysis to assess the developmental progression of anatomical positioning of prefrontal cortex to striatal projections. Corticostriatal axonal territories established at P7 expand in concert with striatal growth but remain largely unchanged in positioning through adulthood, indicating they are generated by directed, targeted growth and not modified extensively by postnatal experience. Consistent with these findings, corticostriatal synaptogenesis increased steadily from P7 to P56, with no evidence for widescale pruning. As corticostriatal synapse density increased over late postnatal ages, the strength of evoked PFC input onto dorsomedial striatal projection neurons also increased, but spontaneous glutamatergic synaptic activity was stable. Based on its pattern of expression, we asked whether the adhesion protein, Cdh8, influenced this progression. In mice lacking Cdh8 in PFC corticostriatal projection neurons, axon terminal fields in dorsal striatum shifted ventrally. Corticostriatal synaptogenesis was unimpeded, but spontaneous EPSC frequency declined and mice failed to learn to associate an action with an outcome. Collectively these findings show that corticostriatal axons grow to their target zone and are restrained from an early age, do not undergo postnatal synapse pruning as the most dominant models predict, and that a relatively modest shift in terminal arbor positioning and synapse function has an outsized, negative impact on corticostriatal-dependent behavior.

## Introduction

Pyramidal neurons in mature prefrontal cortex (PFC) project to spiny projection neurons (SPNs) in the dorsomedial striatum as part of a highly adaptive, recurrent circuit required for reinforcement learning and motivated behaviors^1,2^. Consistent with this role, altered corticostriatal connectivity is a key contributor to human pathological symptoms observed in a wide range of disorders and diseases ranging from schizophrenia and autism to Huntington’s and Parkinson’s^3–5^. Few studies, however, have examined how this pathway develops, and the absence of this knowledge makes it difficult to distinguish normal from pathological innervation patterns, to quantify the impact of molecular factors hypothesized to regulate its development, or to measure the effects of strategies designed to ameliorate developmentally seeded pathologies.

Early studies of developing neural connectivity emphasized broad principles by which topographically organized maps of the sensory periphery distributed across sensory areas, where topographic order is generated in part by regional gradients in neuronal differentiation such that the oldest neurons of one region project to the oldest of another^6^. These ordered connections are then further refined or pruned by ongoing sensory experience-driven activity over clearly defined periods in early postnatal life. Although this framework is well-suited to describing the development of topographically organized unimodal sensory regions, it is inadequate for polymodal cortical and subcortical regions that receive multiple concurrently developing projections from functionally diverse brain areas. Connections from the PFC to the striatum exemplify one such complex pathway. It is with this in mind that we sought to develop an analysis strategy that can assess and compare innervation patterns, suited to defining a loose and potentially flexible topography distinct from the precisely defined maps of the sensory periphery that are generated in sensory areas.

Here, we used a novel approach based on Hotspot Analysis^7^ to define the postnatal developmental progression of PFC innervation of dorsal striatum in mice. Anatomical data were then integrated with synaptic, molecular, and functional measures, and these findings on neurotypical development were used as a basis for characterizing the impact of deleting Cadherin 8 (Cdh8), a Type II cadherin known to contribute to regional, laminar, and cellular identity and that has been hypothesized to contribute to the targeting of PFC to striatal connections^8–11^. The data support a directed growth model where PFC axons are targeted correctly to dorsomedial striatal territories from the earliest stages examined, failing to undergo a developmental period of overgrowth, pruning, and refinement. In the absence of presynaptic Cdh8, PFC-striatal territories shift ventrally, and mice fail to acquire an instrumental learning task, suggesting even modest shifts in anatomical positioning may be magnified behaviorally as information is processed in recurrent loops.

## Results

### Developing PFC corticostriatal axon innervation follows a directed growth model

Corticostriatal projection neurons are generated embryonically between E12.5 and E16.5, and striatal neurons between E12.5 and P2^6,12–14^, but corticostriatal axons wait until after birth (P3-P4) to invade the striatum^15^. To characterize the emergence of PFC terminal axon territories in the striatum, we placed injections of anterograde tracer AAV1.hSyn-TurboRFP into the PFC of mice (Fig. 1B) at P0, P14, and P49, and analyzed projections to the ipsilateral dorsal striatum seven days later (at P7, P21 and P56; Fig. 1A.1-F). Injections were targeted to medial areas of the PFC, including infralimbic cortex (IL), prelimbic cortex (PL), and dorsal anterior cingulate area (ACC) based on functional characterizations of these regions in regulating goal-directed behaviors^16,17^.

**Figure 1.**
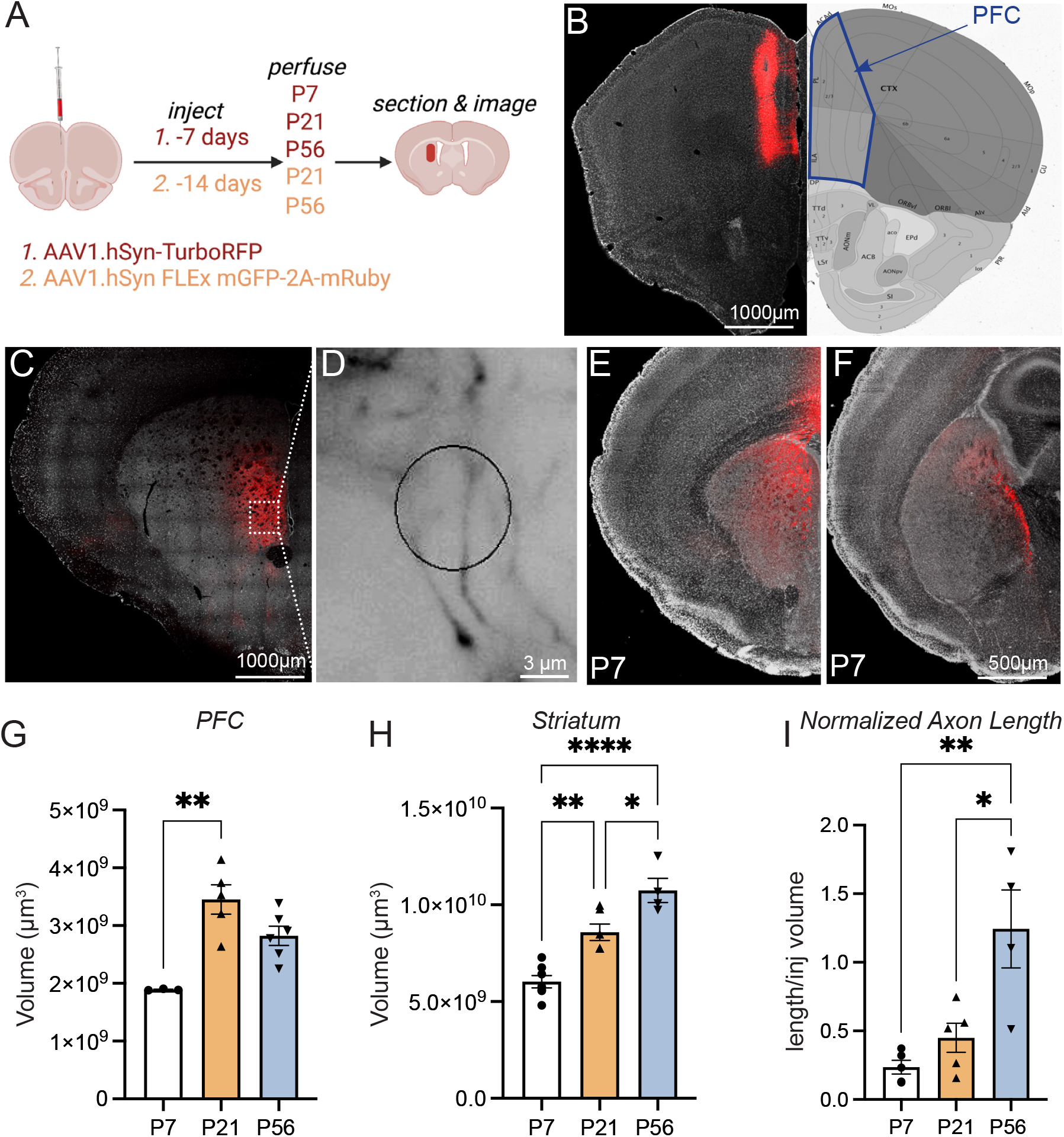
PFC corticostriatal terminal axon extension into dorsal striatum. A. Experiment overview (generated using BioRender). B. Tiled, confocal image of DAPI stained section (white) showing injection site (red) in P56 brain adjacent to image from Allen Brain Atlas showing the region we defined as PFC (ACC + PL + IL, blue line). Examples of PFC axon terminal field (red) in dorsal striatum at P56 captured at low magnification using confocal (C) and at high magnification on a widefield microscope with the Spaceballs circular probe (D, labeling appears in black). Tiled confocal images from P7 brain showing PFC labeled axons (red) in rostral (E) and caudal (F) striatum. Scatterplot/bar graph (G) of PFC volume over time evaluated using Calvalieri estimator (ANOVA; p<0.0028; Tukey’s **p=0.0011). Similar graph (H) tracks striatal volume assessed with planimetry (ANOVA; p<0.0001; Tukey’s: *p = 0.132; **p=0.0012; ***p<0.0001). Total axon length (I) in dorsal striatum normalized to size of PFC injection (ANOVA p=0.0025, Tukey’s: **P7 vs. P56= 0.0025; *P21 vs. P56 = 0.0122).

At P7, axon density was highest rostrally, but labeled axons extended the full rostro-caudal extent of the striatum and were readily detected at the level of the hippocampus (Fig. 1E, F). To capture the extent of axon innervation in relation to the growing striatum, the Cavalieri estimator was used to measure volumes of PFC, striatum, and injection sites (Fig 1G, H), and a systematically applied, stereological “spaceballs” probe^18^ (Fig. 1C, D) was used to estimate total axon length within dorsal striatum (Fig. 1I). The data show that PFC growth was complete by P21 (Fig. 1G), consistent with data in rats^19^, while the striatum continued to grow past the third postnatal week (Fig. 1H). Axon length (normalized to injection site volume) advanced primarily between P21 and P56 (Fig. 1I).

Together, these data show that PFC corticostriatal axons are present throughout the rostral-caudal extent of dorsal striatum as early as P7 but continue to elaborate their terminal length contemporaneously with the increasing volume of the striatum.

In mature mice, PFC axon projections to the striatum cluster dorsomedially^20^. We next addressed whether this pattern results from overgrowth-and-pruning or by directed growth. To visualize and quantify the topographic distribution of PFC axons within the dorsal striatum and to compare the establishment of adult-like territorial patterns across animals and ages, we generated a Python-based clustering algorithm grounded in Hotspot Analysis (Fig. 2). This approach is used in the geosciences and elsewhere to identify significantly clustered, spatially discrete data^7^. Following targeted cortical injections of anterogradely-transported fluorescent proteins (Fig. 2A), tiled images were captured through the rostral-caudal extent of the striatum following the sampling strategy used for stereology (see Methods). In hemi-sections, bundles of axons in the internal capsule were masked and removed from analysis based on their intensity (Fig. 2B,C); Hotspot Analysis was then applied to the remaining axon terminal labeling patterns within the striatal neuropil. The pipeline examined 20 × 20 pixel regions, or “neighborhoods”, and assigned values based on z-scores relative to the same image in which all pixel intensities were scrambled (Fig 2C, G “Image” vs “Randomized”). Thus, each image served as its own control. Positive scores reflect “hot spots” and negative scores, “cold spots”. The data were visualized by *(1)* fluorescence intensity-based heat maps (Fig. 2H, K); *(2)* mountain plots showing the distribution of z-scores along the dorsal-ventral and medial-lateral axes of the striatum (Fig. 2H, K); and *(3)* histograms where Z-scores were subdivided into four topographic quadrants, dorsomedial (DM), dorsolateral (DL), ventromedial (VM), ventrolateral (VL) (Fig. 2I, L), based on the centroid of each region of interest (ROI), which was defined by the contours of the striatum; Fig. 2G, J).

**Figure 2.**
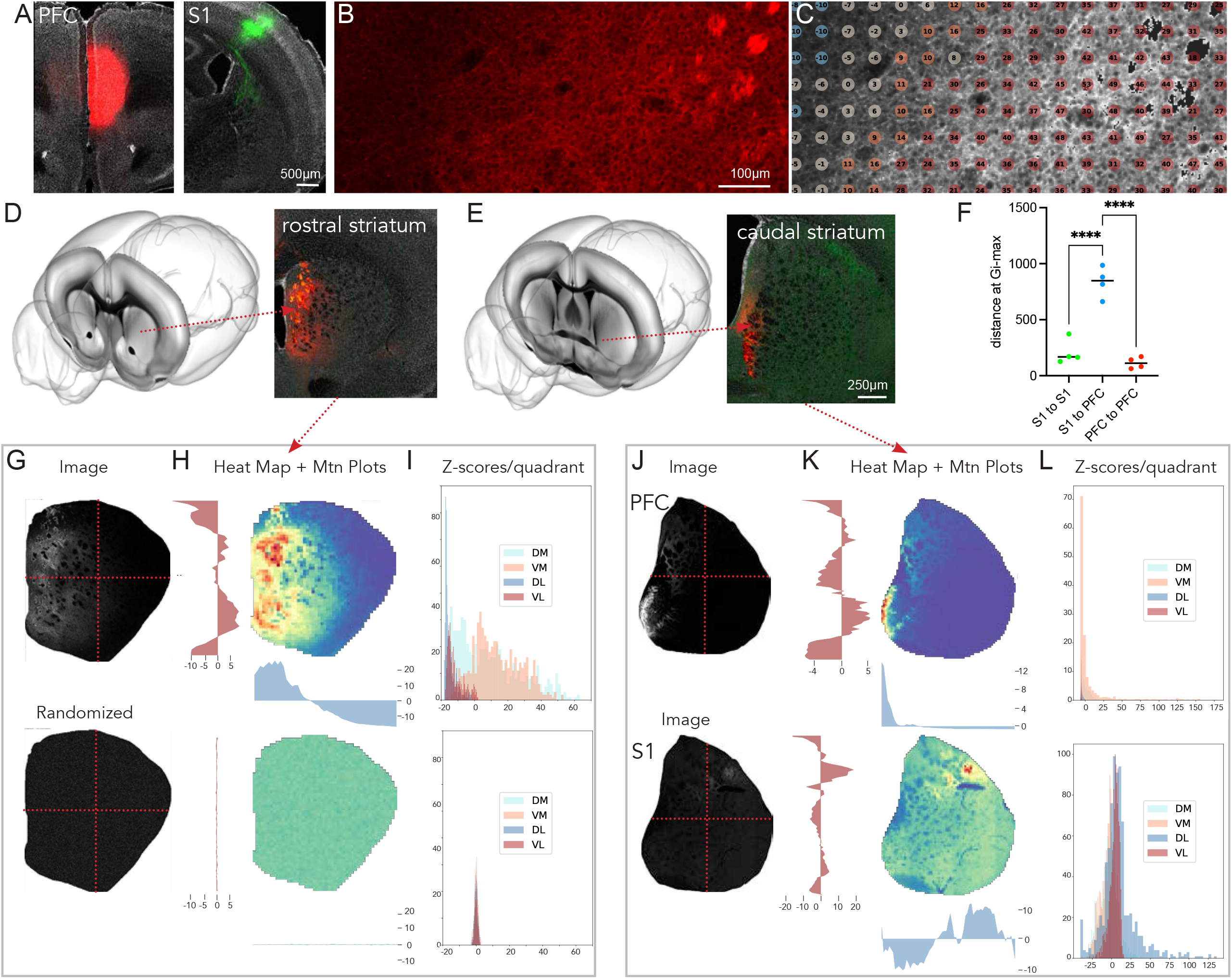
Hotspot Analysis for comparing PFC axon territories. A. Tiled confocal images of sections through adult mouse brain showing injection sites for AAV-mRuby in PFC (A) and AAV-GFP in S1 stained with DAPI (white). Tiled confocal image (B) through rostral striatum showing fine, axon arbors (red) as well as axon fascicles. Same image (C) with axons displayed in white, fascicles eliminated by thresholding (black) and z-score map from Hotspot Analysis. Warmer colors have higher z-scores. Images through rostral (D) and caudal (E) striatum showing terminal labeling patterns for PFC and S1 axons. Approximate rostral-caudal levels are depicted in cartoons accessed from MouseLight https://registry.opendata.aws/janelia-mouselight^68^. Scatter plot (F) compares the XY distance between neighborhoods having the highest z-scores following injections into the cortical sites indicated on the x-axis. (One way ANOVA, p<0.0001; ****Tukey’s, p<0.0001). G-I show outputs from Hotspot Analysis pipeline (see also text) from section in D with upper panels displaying data from the actual image and lower panels, from the same image in which pixels have been randomized. J-L show Hotspot outputs from section in E with PFC terminal arbors at the top and S1 terminal arboras at the bottom. Quadrants used in I and L are marked by the red dotted lines seen in G and J. Abbreviations: Mtn, mountain; DM, dorsomedial; VM, ventromedial; DL, dorsolateral; VL, ventrolateral. S1, somatosensory cortex area 1.

To first validate the approach, we injected AAVs expressing different fluorescent proteins into PFC (turboRFP) and primary somatosensory cortex (S1; mGFP) (Fig. 2A) of adult mice. We picked these two cortical regions because striatal terminal axon territories from these sites should be largely distinct and non-overlapping^20,21^ (Fig. 2D, E). Hotspot Analysis showed that peak z-scores corresponding to PFC or S1 terminal fields lie within completely separate striatal territories (Fig. 2G-L) and occupy zones that are consistent with published work that has described these terminal projections in adult mice^20,21^. In the examples shown, a section through the rostral striatum shows only PFC axons, and these are clustered in a medial strip (Fig. 2D, H, I); in a section through the caudal striatum, PFC axons cluster ventromedially and S1 axons cluster dorsolaterally, differences that can be readily appreciated in the mountain plots as well as in the quadrant-based histograms, where positive z-scores lie within completely different quadrants (Fig. 2E, K, L). To compare the positions of terminal fields across animals, we took the XY coordinates of the neighborhood in which the Gi statistic (a ratio of the total pixel values in a neighborhood to the global total, see Methods) was maximal (or having the highest Z-score) and then measured the distances between coordinates in each mouse. The data show that maximum Gi is observed at similar positions for either S1 or PFC injections across animals, that distances between S1 and PFC sites are greater, as expected, and also that they are consistent across animals (Fig. 2F).

We next used Hotspot Analysis to define and compare the striatal territories occupied by PFC afferents as they develop. At P7, in the rostral striatum, clustered PFC terminal axons occupy much of the medial two-thirds of the striatum, a distribution that is clear in the medial to lateral mountain plots and from the dominant hot-spots that occupy the DM quadrant and the significant cold-spots that occupy the DL quadrant (Fig. 3A). Moving caudally, PFC terminal axons become contracted into a more restricted, apostrophe-shaped region, with a wider, circular zone sitting dorsomedially and a narrower tail extending ventromedially; both zones show as positive deflections in the mountain plots and are evident in the quadrant distributions, which show ventromedial and dorsomedial hotspots (Fig. 3A, B). The dorsal rim of the striatum and much of the lateral striatum show negative z-scored cold spots (Fig. 3A-C). This basic pattern, evident at P7, is remarkably similar to that observed at P21 and P56 (Fig. 3G, H). Variations in the patterning could be tracked to slight differences in injection site placement rather than to age. For example, cortical injections sites that included the ACC produced more central striatal terminal axon labeling (the rounded portion of the apostrophe), while cortical injections that extended ventrally to include dorsal peduncular cortex (DP) yielded more ventral terminal axon labeling (Fig. 3D, E), a difference that could be detected by an increase in the distance between maximal Gi scores (Fig. 3F). Collectively these data indicate that the topography of striatal territories occupied by PFC axons is largely established by P7 and changes little with subsequent age, thus appearing to be generated prospectively through directed growth.

**Figure 3.**
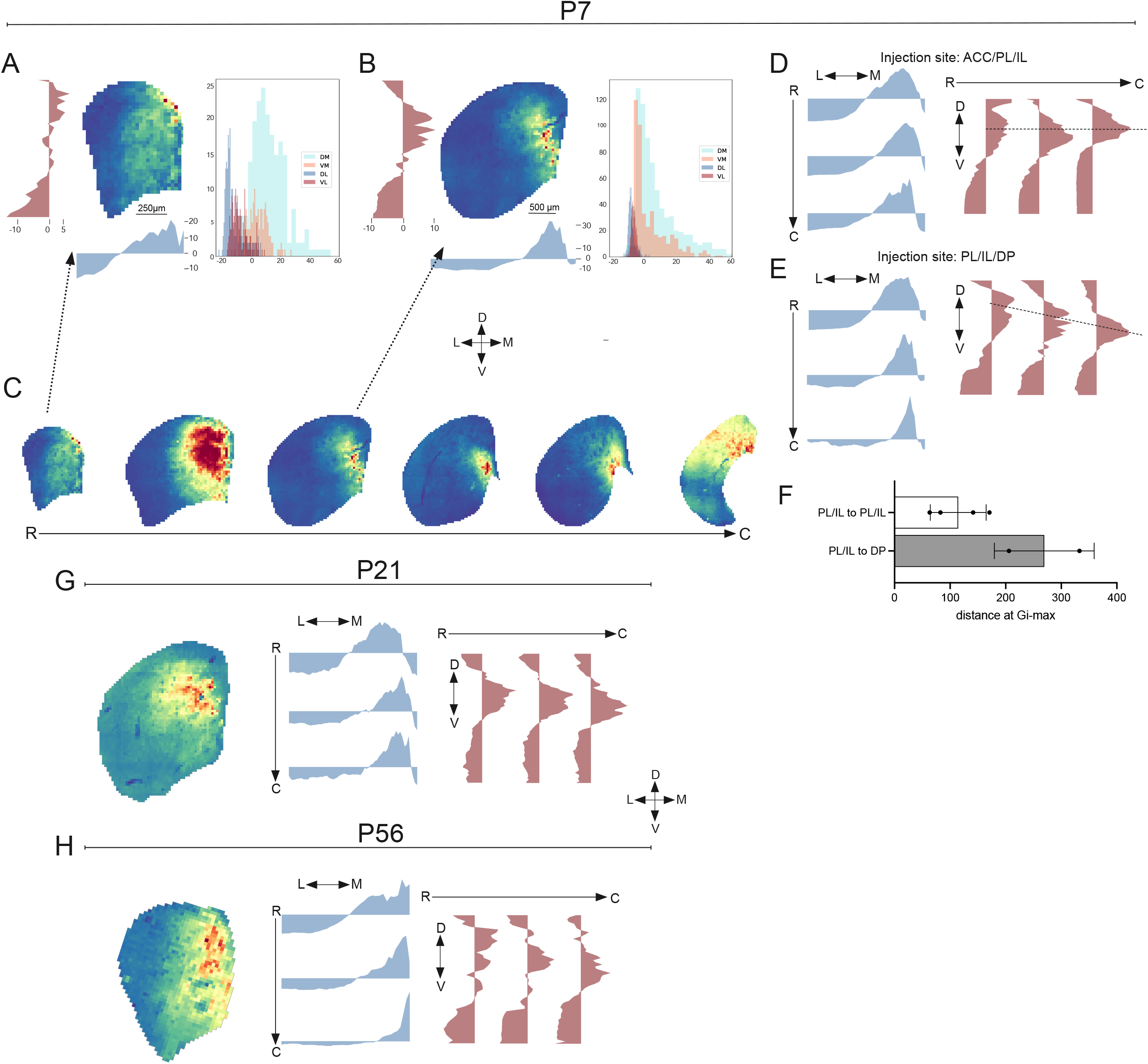
PFC terminal axon territories in striatum are established by P7. Outputs from Hotspot Analysis at rostral (A) and midstriatal (B) levels showing heatmaps, mountain plots and quadrant maps at P7. Heat maps running rostral to caudal are shown in C. Mountain plots generated from an injection site that included ACC (in addition to PL and IL) showed consistent medial/lateral and dorsal/ventral peaks, moving caudally (D; dotted black line on D/V plots), but injection sites including DP (dorsal peduncular area) (E) produced peaks that move ventrally at caudal levels, consistent with a greater XY distance between peak z-score neighborhoods produced by PL injections to those from injections including DP shown in scatter/bar plot (F). At P21 (G) and P56 (H) the Hotspot outputs appear very similar to what is observed at P7. Abbreviations as in Figure 2 or in text.

### Corticostriatal synaptogenesis extends through postnatal development

Corticostriatal synapse densities were estimated by quantifying appositions of presynaptic VGLUT1- and postsynaptic Homer1-immunolabeled puncta imaged at high magnification (Fig. 4A, Methods). VGLUT1 is expressed in all axon terminals from the cortex, amygdala and hippocampus, but the latter two sources account for only a small fraction of the total^22,23^ (Fig. 4A; Methods). Homer1 is common to most excitatory postsynaptic densities. At P7, apposition density in the lateral striatum (Region B) was significantly higher than that observed either dorsally (Region A) or ventrally (Region C) (Fig. 4B-D), perhaps reflecting its earlier generation^12,24^, but by P21 this regional difference was no longer evident (Fig. 4C, D). This suggests that the ingrowth of sensorimotor inputs laterally precedes PFC inputs medially. Mean apposition density across the striatum did not change between P7 and P21 (Fig. 4H), suggesting that the pace of synaptogenesis during this time frame was sufficient only to match the gains in striatal volume. This early period was followed by a surge in apposition density between P21 and P56 (Fig. 4H). Although the density of VGLUT1 terminals increased linearly over time (Fig. 4F), the rise in Homer1 puncta density coincided with apposition density (Fig. 4G), suggesting that postsynaptic maturation was rate limiting. Sizes of VGLUT1 puncta were larger at P7 than at the later ages, suggesting that vesicles in young boutons could be more numerous or (more likely^25^) loosely packed (Fig. 4E).

**Figure 4.**
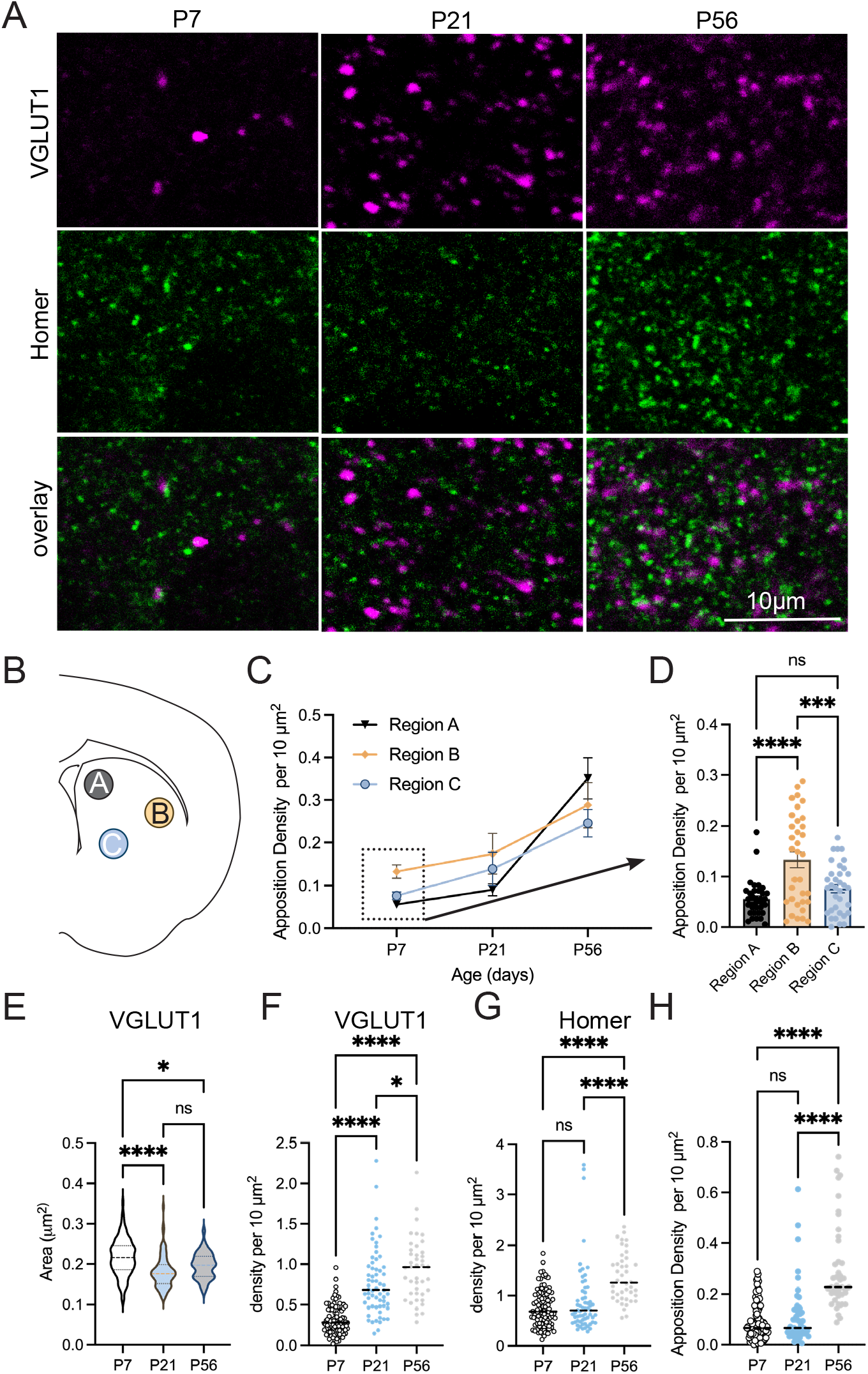
Synapse density increases steadily. Images of VGLUT1 (magenta) and Homer (green) immunolabeling in sections through dorsal striatum and captured on a confocal microscope (A). Diagram in (B) shows the regions from which images were captured (A: gray, dorsal); B: beige, lateral; C: blue, ventral). Line graph in (C) plots VGLUT1 X Homer apposition density (see methods) for each region examined and bar/scatterplots in (D) compare regional densities at P7. Violin plot in E compares area of VGLUT1 terminals, and scatter plots in F, G and H, compare densities of VGLUT1 terminals, Homer1 puncta and appositions over time (as shown). (E-H) White (P7); blue (P21); gray (P56)

Whole-cell patch-clamp recordings in acute striatal slices were used to compare the functional properties of dorsomedial SPNs at P21 and P56. Intrinsic membrane excitability changed significantly over this period. Rheobase--the minimal current needed to generate an action potential--increased, while membrane resistance decreased between P21 and P56 (Fig. 5A, B). Excitatory synaptic properties (frequency and amplitude), however, were unchanged. Spontaneous EPSCs (sEPSC) displayed similar mean interevent intervals and current amplitudes at the two ages (Fig. 5C, D), an unexpected finding in light of the rise in synapse (apposition) density between P21 and P56 (Fig. 4H). We reasoned this could be because EPSCs reflect glutamatergic sources (thalamic, amygdala, ventral hippocampus) outside of cortex that could have a strong functional impact despite their small number. Thus, to compare the strength of identified PFC-dorsomedial striatal synapses at these two ages, we used an optogenetic approach. AAV was used to express channelrhodopsin (CaMKIIa- hChR2(H134R)-eYFP) in PFC striatal projection neurons, and their terminals in dorsomedial striatum were optically stimulated in acute slices with increasing light intensity. The input-output relationship of optically-evoked synaptic responses was evaluated by whole-cell recording from SPNs in the dorsomedial striatum (Fig. 5E). There was a substantial and significant increase in optical-EPSC (oEPSC) amplitude at P56 compared to P21 (Fig. 5F). These findings support that synaptic connectivity specifically between PFC and striatum becomes stronger between P21 and P56.

**Figure 5.**
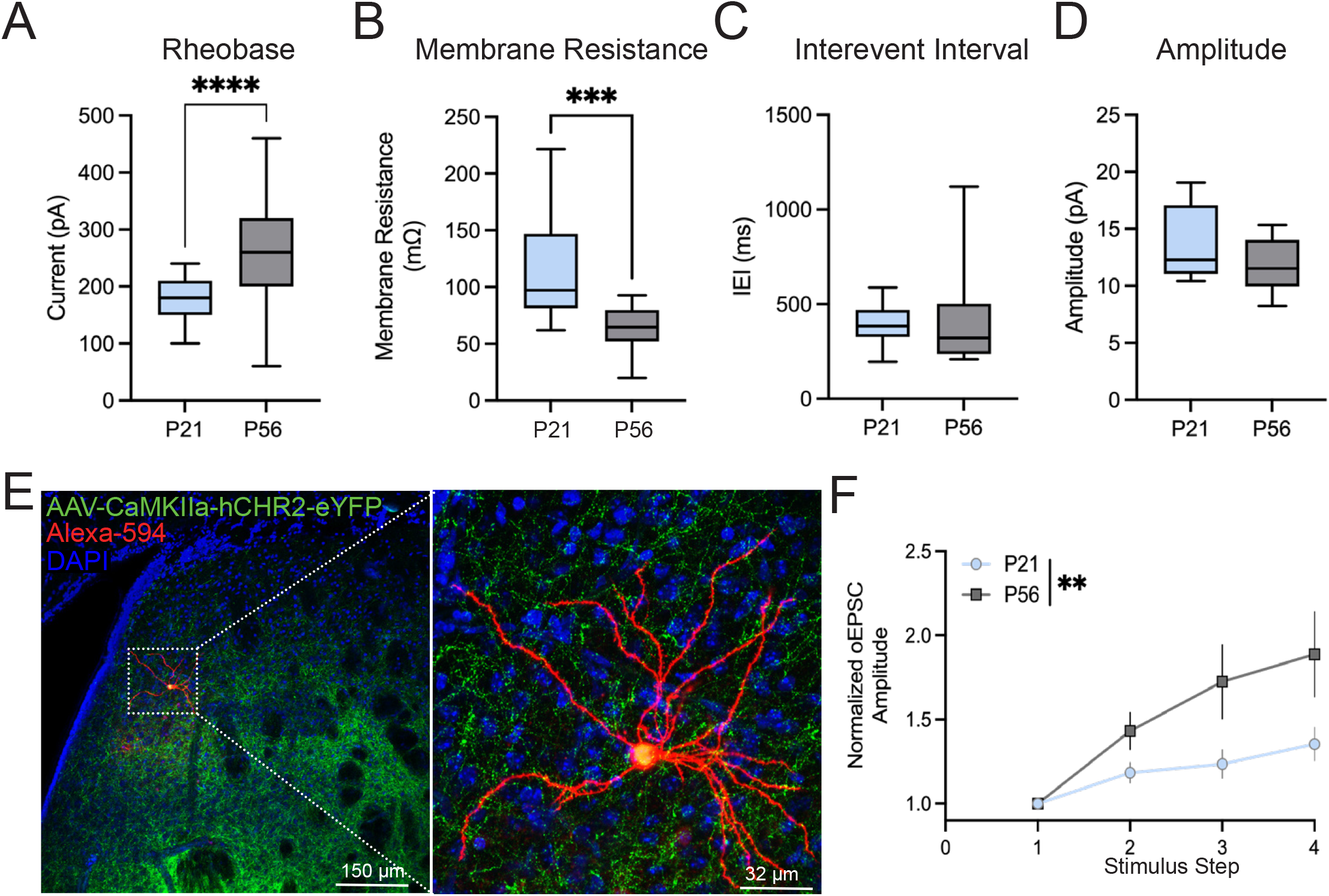
SPN intrinsic properties change and PFC synapses strengthen over time. Scatter/bar plots of whole-cell recording data comparing intrinsic properties: rheobase (A; p = 0.0001, t-test) and membrane resistance (B; p = 0.0007, Welch’s t-test; P21:17 cells/6mice; P56: 19 cells/6 mice); and synaptic properties, interevent interval (C; p = 0.4875; Mann-Whitney) and amplitude (D; p = 0.1643; Mann-Whitney; P21: 16 cells/6 mice; P56: 18 cells/6 mice) at P21 (blue) and P56 (gray). Confocal images in E show the PFC expression of AAV1-CaMKIIa-hChR2(H134R)-eYFP (green) in medial striatum following injection in PFC. Alexa594-labeled SPN (red; filled during whole-cell recording) is shown at higher magnification at right; DAPI (blue). Line graph in F shows output amplitude normalized to the first detected response (y-axis) in relation to input stimulus step (x-axis) at P21 and P56. Data were compared using a Mixed effects analysis (n=8 cells/ 5 mice at P21 and n=13 cells/5 mice at P56; Age effect, 0.0052, Intensity effect, 0.0011).

### Presynaptic cadherin 8 is largely dispensable for targeting PFC axons to striatal territories

Expression of the cell adhesion protein Cdh8 is particularly enriched in PFC and dorsal striatum as PFC axons are invading and establishing their territories^10,11^. In the retina, Cdh8 plays an instructional role in targeting bipolar cell axon arbors^8^, and we wondered whether it might play a similar role in organizing striatal axon terminal territories. To test this, we generated a conditional, *loxP*-dependent *Cdh8*-knockout mouse using a construct that also contained a *Frt*-flanked LacZ reporter (Fig. 6A; Methods). Prior to deleting the reporter cassette, we used LacZ expression to confirm that the construct was inserted appropriately and that Cdh8 is enriched in ACC, PL, IL, and striatum^10^. Consistent with prior work, Cdh8 expression shows a medial (high) to lateral (low) gradient in cortex and a modest, dorsal (high) to ventral (low) gradient in dorsal striatum with little expression evident in ventral striatum (Fig. 6B). To test the efficiency of *loxP*-mediated recombination, *Cdh8*^flfl^ mice were crossed with *Nex(NeuroD6)-Cre*^+/-^ mice, in which Cre recombinase is expressed by excitatory neocortical neurons but not by striatal neurons^26^ (Fig. 6C). Western blots of regional dissections taken from littermates show near complete loss of Cdh8 in PFC (the remainder is consistent with low levels of Cdh8 reported in inhibitory neurons^27^), but unchanged Cdh8 expression levels in striatum as expected (Fig. 6D). Blots from the same tissue also show that levels of N-cadherin and the universal cadherin binding partner, ß-catenin, were unchanged by Cdh8 deletion, suggesting expression-based compensation by other cadherins is unlikely (Fig. 6C).

**Figure 6.**
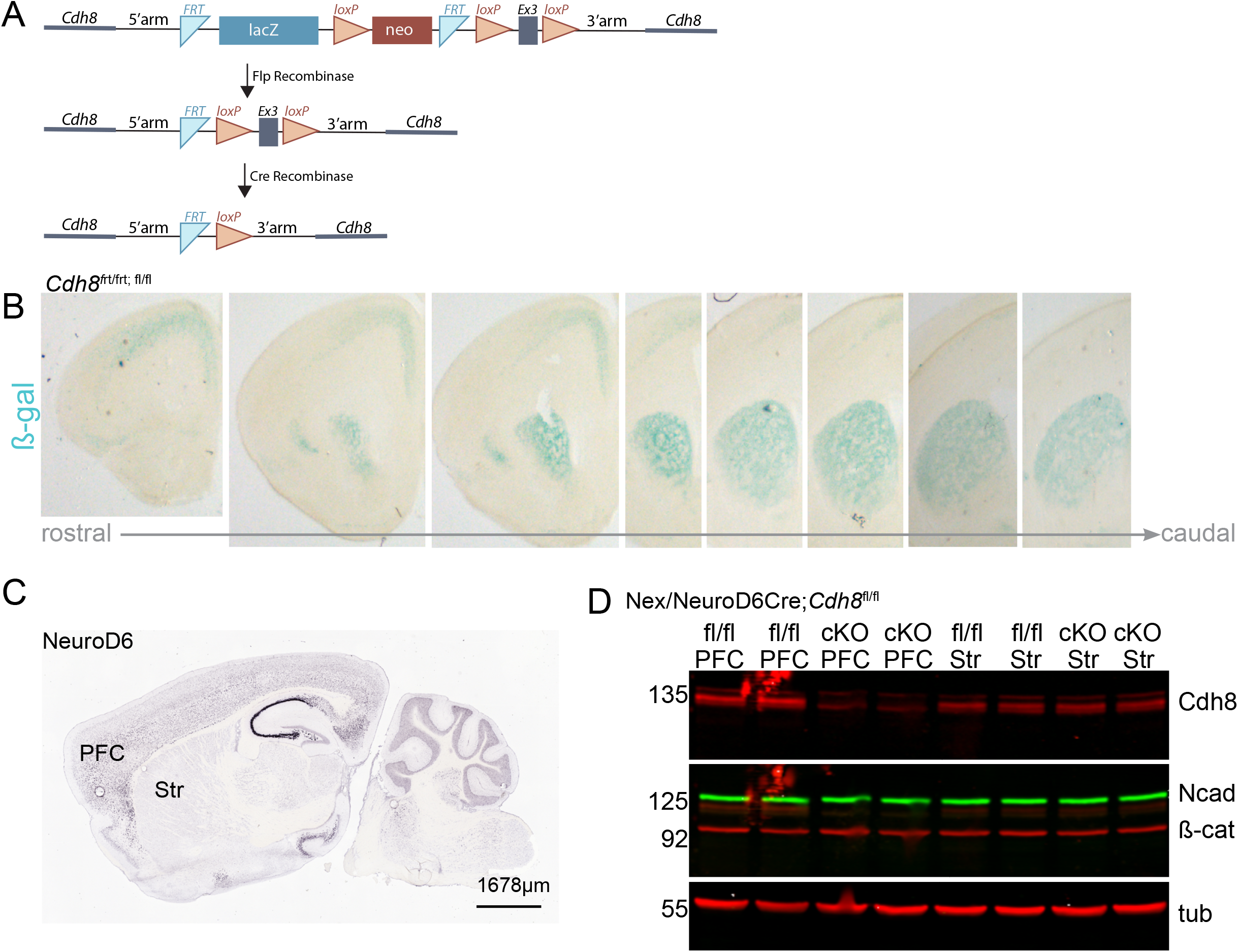
Generating a Cdh8^fl/fl^ mouse. Diagram (A) shows the targeting construct obtained from the Knockout Mouse Project (KOMP) and alleles generated following Flp and Cre-mediated recombination in mice. Rostral to caudal sections taken on a macroscope (B) show LacZ staining in sections taken from the Cdh8^frt/frt; fl/fl^ mouse. In situ hybridization data (C) show Nex/NeuroD6 mRNA expression in PFC and none in striatum (Str; image reproduced from the Allen Brain Atlas; ^32^. Western blots (D) show Cdh8, N-cadherin (Ncad), ß-catenin (ß-cat) and tubulin (tub) levels in lysates taken from the PFC or striatum in Cdh8^fl/fl^ (fl/fl) control compared to Nex Cre; Cdh8^fl/fl^ (cKO).

We next generated a line of conditional Cdh8 knockout mice (Cdh8-cKO) by crossing *Cdh8*^fl/fl^ mice with *Rbp4*-*Cre*^*+/-*^ expressing mice in order to conditionally disrupt Cdh8 expression in corticostriatal projection neurons^28–31^. Adult Cdh8-cKO mice showed no gross anatomical abnormalities by Nissl-staining, and when crossed to a tdTomato-Cre reporter line (Ai9), sections through the forebrain showed that Cdh8-cKO neurons migrated to their appropriate laminar positions and their axons projected normally to the striatum in a matrix-like pattern, similar to *Rbp4*-*Cre*^*+/-*^ control mice^30,32^ (Fig. 7A, D). At the same time, Hotspot Analysis revealed modest differences in innervation patterns, with Cdh8-cKO axons shifting ventrally and medially relative to controls (Fig. 7B, C, E, F).

**Figure 7.**
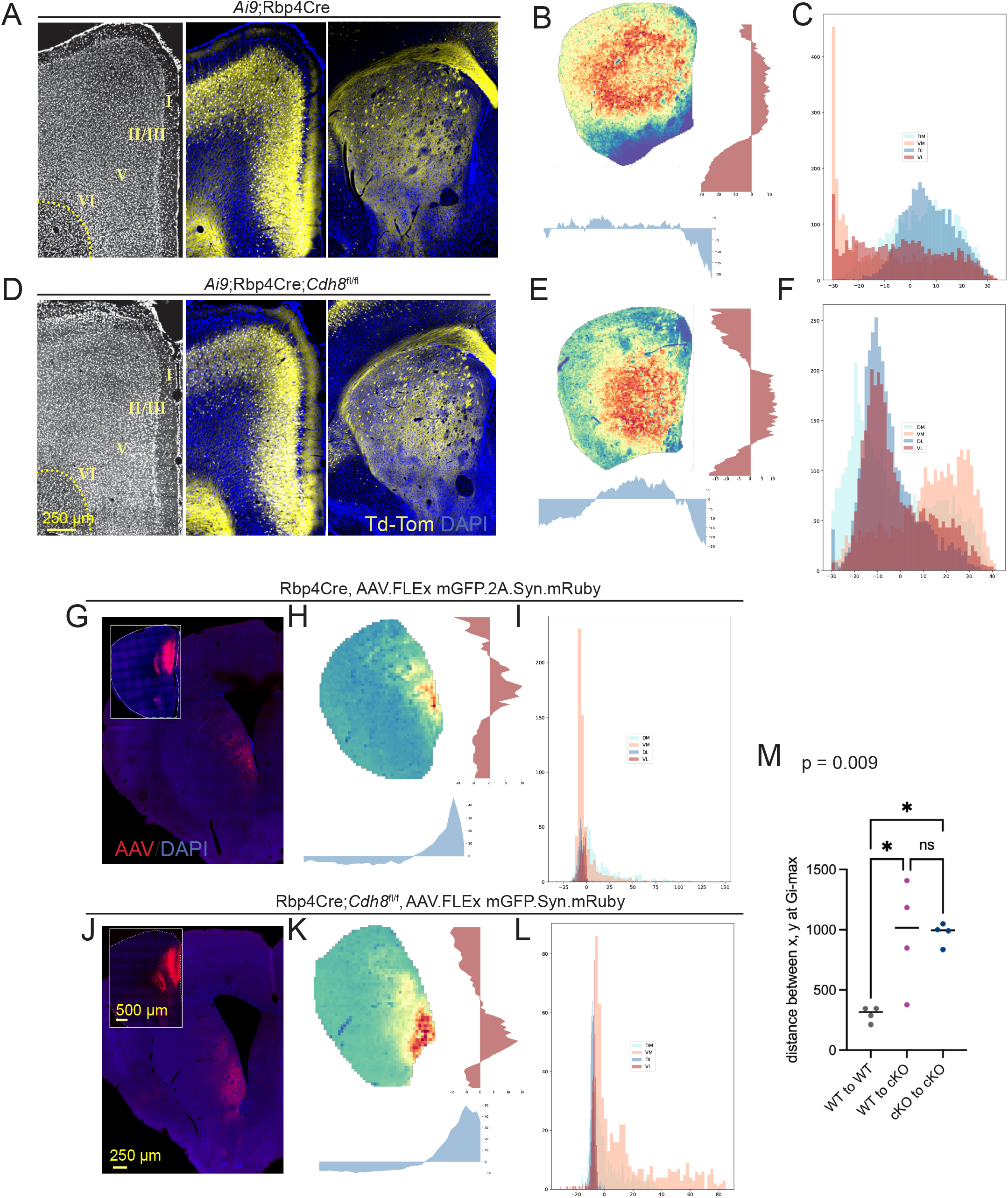
Presynaptic Cdh8-cKO shifts striatal territories ventrally. A-F show data obtained from Rbp4- Cre control (A-C) and conditional Rbp4Cre;Cdh8^fl/fl^ knockout tissue (D-F) from P21 mice crossed to a tdTomato Cre-reporter line (Ai9). Tiled confocal images in A and D show prefrontal cortex (left: DAPI, white and middle: DAPI, blue, tdTomato, yellow), and dorsal striatum (right). Cortical layers indicated by roman numerals and magnification shown in D. Heatmaps, mountain plots and quadrant plots from control (B, C) and Cdh8-cKO (E, F), reveal a modest ventral shift in corticostriatal territories. G-M shows tiled confocal images (G, J) and corresponding hotspot output in mid-striatal level sections (H, I; K, L) obtained from P56 Rbp4Cre control (G-I) and conditional Rbp4Cre; Cdh8fl/fl knockout (J-L) tissue from mice following injections of Cre dependent AAV.mGFP-2A-Syn.mRuby in PFC (insets in G, J). AAV expression in red; reflects mGFP at the injection site and Syn-mRuby in the striatum; DAPI in blue; respective magnifications shown in J. Dot plot (M) compares distances between mice of X, Y coordinates at site of Gi-max. 1-way ANOVA, p = 0.009; Tukey’s multiple comparison test, *p<0.017.

To distinguish and compare the impact of Cdh8 disruption on PFC axons specifically, a Cre-dependent virus (AAV1-hSyn-mGFP-2A-synaptophysin-mRuby) was injected into PFC of Cdh8-cKO and control mice and striatal terminal fields were compared using Hotspot Analysis. Heatmaps and mountain plots show that PFC terminal axon territories exhibit a modest ventral shift (Fig. 7G-L and Suppl. Fig. 1 A-C). This was most evident at mid-striatal levels, with Cdh8-cKO axons more robustly invading ventrolateral territories and avoiding dorsomedial zones (Fig. 7G-I vs. J-L; Suppl. Fig. 1 A-D).

Maximum Gi neighborhoods were observed at similar positions across control mice, but consistent with targeting defects, the distances were greater between control and Cdh8-cKO mice as well as between Cdh8-cKO mice (Fig. 7M). These data support that in the absence of presynaptic Cdh8, positioning of PFC terminal axon territory shifts and is more variable than controls.

### Presynaptic reduction in Cdh8 expression has modest impact on synapse function

Cdh8 is also found concentrated at asymmetric synapses in dorsal striatum^10^. To test whether presynaptic deletion of Cdh8 regulates synaptogenesis, we compared VGLUT1/Homer1 apposition density in dorsal striatum in Cdh8-cKO and control (Rbp4Cre and wildtype) mice. No differences between genotypes were observed at either P21 or P56 (Fig. 8A-C). Comparisons across striatal regions as defined in Figure 4B, also showed no impact of genotype (Suppl. Fig. 2A, B). In contrast, whole cell recordings showed a significant increase in sEPSC interevent intervals in SPNs from Cdh8 cKO mice compared to littermate controls at P21 that were sustained at P56 (Fig. 8D, F, G). A significant decrease in amplitude was also observed at P21 but its small magnitude and absence at P56 suggests it is unlikely to be biologically meaningful (Fig. 8E, H). Although we observed no changes in glutamatergic appositions, cadherins can influence dendritic spine shape in hippocampus^33,34^ and decreased afferent activity during striatal development might be anticipated to decrease protrusion density or slow maturation^30^. To examine these possibilities, we labeled biocytin-filled SPNs with Alexa594, imaged them at high magnification, and analyzed all protrusions using NeuronStudio (Fig 8I, Suppl. Fig. 2C). Protrusion density increased between P21 and P56 as expected, but there were no genotype-dependent differences detected at either age (Fig. 8J). There were also no significant differences in protrusion head diameter, but a modest increase in protrusion length was observed at P56 (Suppl. Fig. 2C-E). Collectively the data suggest that Cdh8 has a modest impact on synapse maturation but a larger effect on glutamatergic synapse function.

**Figure 8.**
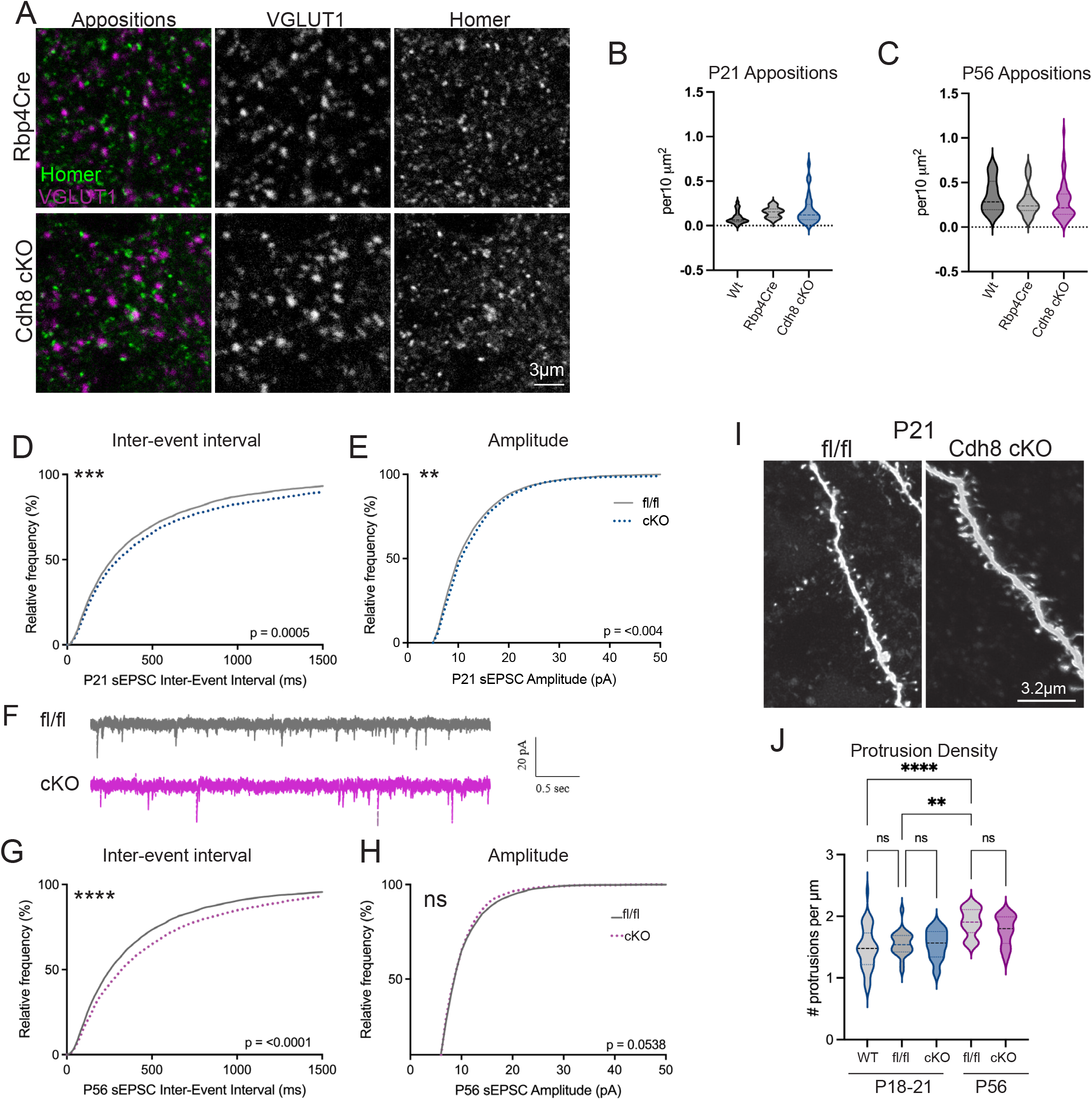
Cdh8 cKO reduces presynaptic function. Confocal images of sections through striatum of P56 control (Rbp4Cre) and Cdh8 cKO mice immunolabeled for VGLUT1 and Homer, shown individually and as an overlay (A; as shown). Violin plots in (B) and (C) compare VGLUT1/Homer apposition density in medial striatum assessed using strategy shown in Figure 5 (1-way ANOVA; p>0.05). Cumulative frequency plots and traces from whole-cell patch clamp recordings in SPNs from P21 (D, E) or P56 (F-H) control (Cdh8^fl/fl^) or Cdh8-cKO slices showed significantly longer sEPSC interevent intervals at both ages (H, G, H; Kolmogorov-Smirnov test ***p=0.0005; **** p<0.0001) and a very modest increase in sEPSC amplitude (Kolmogorov-Smirnov test p=0.0043, P21 and p =0.05, P56; n=15-20 cells/ 5 animals per group). Confocal images (maximum intensity projections, deconvolved) of dendritic segments from filled medial striatal projection neurons at P21 (I). Violin plot (J) compares protrusion density at P21 and P56 (1-way ANOVA, p < 0.0001; Tukey’s post test: age- dependent differences P21 vs P56 fl/fl, p=0.0011; P21 wildtype vs. P56 fl/fl, p <0.0001. Within an age, no differences were detected between fl/fl and wildtype (p=0.8388) or between fl/fl and cKO (p= >0.9 at P21; p = 0.7074 at P56).

### Instrumental learning

In a final set of experiments, we tested whether Rbp4Cre-mediated Cdh8 deletion altered action-outcome associations that rely on intact prefrontal-dorsomedial striatal connectivity. Using touchscreen-based operant chambers, we subjected male mice to a 4-day, instrumental learning paradigm in which they learn to associate an action with a reward. In wildtype mice, this instrumental paradigm drives goal-directed learning, substantiated by sensitivity to outcome devaluation ^35,36^. Under a continuous reinforcement (CRF) schedule on day 1, all mice learned to associate a nose poke to a lit screen with a strawberry milk reward, displaying a similar number of responses (Fig. 9A; CRF). On subsequent days 2 - 4 under a random interval (RI) reward schedule, control mice progressively increased number of responses as the interval extended from 15 to 30 seconds, as expected^35,37^ (Fig. 9A). In contrast, Cdh8-cKO mice showed no development of instrumental learning across training sessions (1-way ANOVA, p =0.3; test for a linear trend: p = 0.1785) (Fig. 9A). The failure of the Cdh8- cKO mice to learn this task prevented us from progressing to reward devaluation experiments that could be used to parse goal-directed from habitual responding. There were no significant genotype-dependent differences in overall motor activity (inferred by beam breaks) in any of the sessions (Fig. 9B), nor were there significant differences in mean reward collection latencies (Fig. 9C), suggesting that reduced performance of Cdh8-cKO mice was neither a function of reduced locomotor activity nor reduced motivation for the reward. Additionally, neither locomotion nor reward collection latencies correlated significantly with performance on the instrumental learning task on the first day of RI-30 testing (Spearman, reward collection latency: R=-0.56, p = 0.13; locomotion, R=0.20, p= 0.63), also supporting that the deficits observed in Cdh8-cKO mice are in instrumental learning. The poor performance of the Cdh8-cKO mice suggests that the modest anatomical and physiological differences observed are amplified in the context of a behavior that relies on intact and fully functioning connectivity between PFC and striatum and other sites.

**Figure 9.**
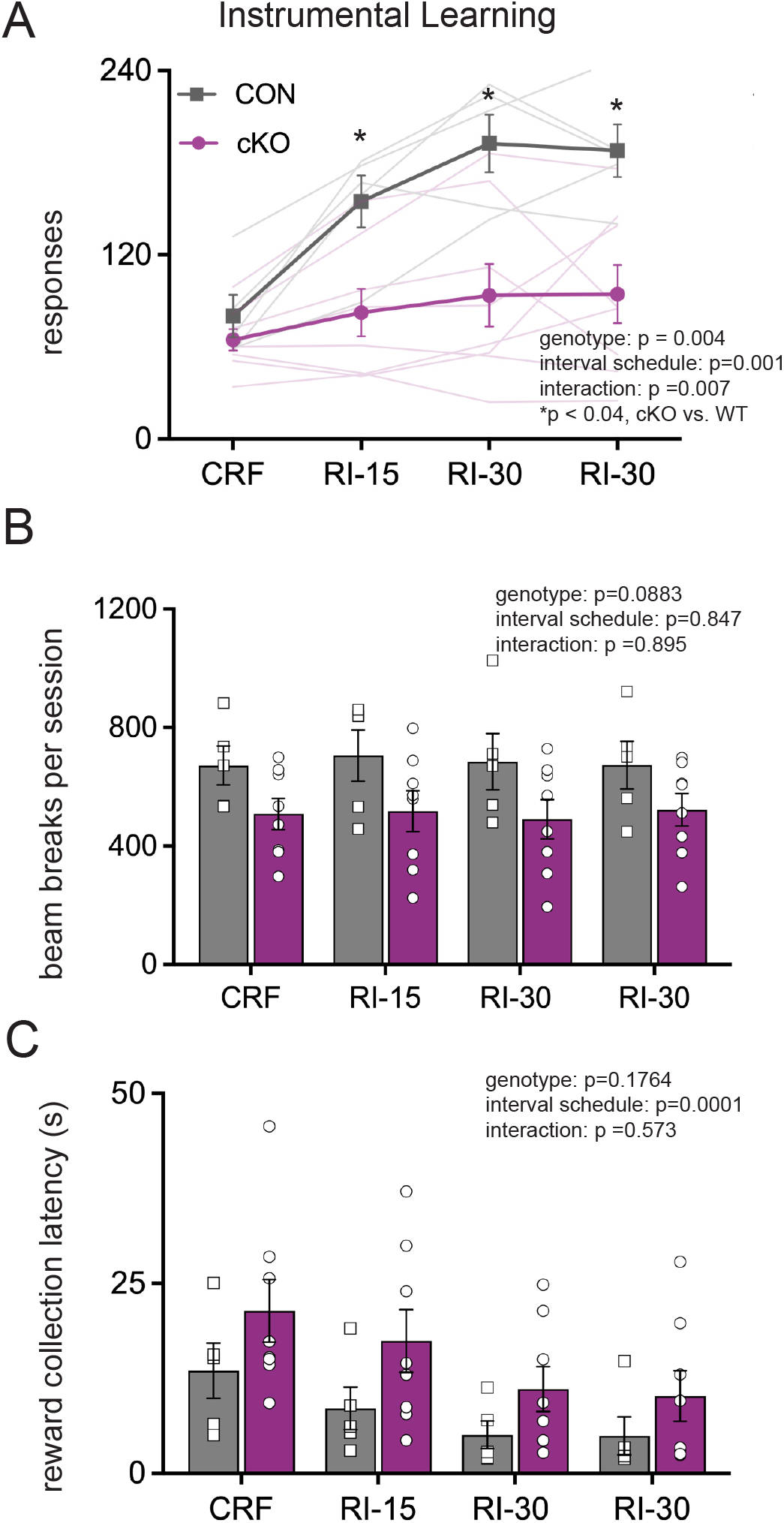
Cdh8-cKO mice exhibit reduced performance in an instrumental learning task. Line graph (A) shows mean response times during an instrumental learning paradigm in which mice were rewarded every time they touched a screen (continuous reinforcement, CRF); or at random intervals (RI) within 15 or 30 second windows, as indicated (bars are SEM; n=5 wildtype control CON; gray and 8 Cdh8 cKO mice; purple). Trace lines show individual animals. Cdh8 cKO mice failed to learn this task (A). Bar graph/scatter plots show beam breaks (B) and reward collection latencies (C) for each session. While there were no genotype dependent differences in collection latencies, both groups showed a decrease over time, supporting that the mice were motivated to obtain the reward. 2-way ANOVA data are reported in each figure

## Discussion

Here we combined circuit-based mapping with application of a novel terminal axon clustering algorithm (Hotspot Analysis) to provide new details on the sequence over a developmental timecourse through which PFC axons establish topographically organized terminal territories within striatum and to show how they are modified by Cdh8. The data show that the adult-like, topographic distribution of PFC projections to dorsomedial striatum is largely defined by P7, with no evidence for widescale axonal or synaptic pruning. Conditional, presynaptic deletion of Cdh8 shifted PFC terminal axon territories ventrally, diminished synapse function, and produced significant deficits in instrumental learning.

One of the key findings here is also the simplest: the adult-like topography of PFC projections to striatum emerge developmentally by a targeted growth mechanism. Although there are noteworthy exceptions (e.g ^38,39^), the literature is dominated by models in which axon growth is promoted and restrained toward a principal target within which terminal arbors overshoot and then are subsequently pruned to their final position/s^40^. During development, PFC axons grow toward and innervate a defined territory and arborize within that territory in a manner that is proportional to striatal growth. The onset of cortical innervation lags behind the generation of striatal projection neurons^15^ suggesting that axons from multiple cortical regions invade the striatum simultaneously such that position of cortical projections is driven in part by the location of invading neighbors. At the same time, however, glutamatergic thalamic innervation precedes cortical innervation^41^. Recent work has shown that in adult mice, the parafascicular nucleus can be divided into dorsal-ventral strips or zones based on thalamostriatal projections and that medial parafascicular nucleus axons are confined to a medial zone of striatum^42^ that cleaves close to the PFC terminal zone, laying out the possibility that they may generate a template for cortical axons to follow.

The data also support that corticostriatal synapses do not undergo a phase of widescale synapse pruning as part of the process of circuit maturation. Synaptogenesis in lateral striatum led medial, consistent with the developmental gradient of SPN generation^6,43^, and there was a steady increase in corticostriatal (VGLUT1-labeled) pre-and postsynaptic appositions and dendritic spine density that dovetail with prior reports of a steady increase in SPN dendritic spine and asymmetric synapse density in dorsal striatum^44–48^. These events are in keeping with the strong influence of emergent cortical activity on striatal synapse development and maturation^30,46^ and the close relationship between the generation of dendritic arbors and synapse formation^45,49^. However, unlike many brain areas^50,51^, there is little evidence here or in previous work that cortical inputs to dorsal striatum undergo a period of significant synapse elimination as part of a process of circuit refinement during development and early adulthood. It may be that use-dependent synapse sculpting is exerted differentially, as may be the case with different cortical layers^52^, or contemporaneously, if increased input from one cortical area is balanced by loss from another, possibilities we would not have detected. At the same time, it should be underscored that corticostriatal synapses *can* be eliminated under pathological circumstances, as happens in adult rodents in response to 6-OHDA lesion^53^ or in humans with Parkinson’s^54^, but such loss in striatum appears to be motivated by circumstances that lie outside the experience or competition-dependent pruning commonly associated with circuit formation during development. The absence of pruning is also consistent with the idea that corticostriatal territories are largely directed from the outset.

Although corticostriatal synapse density rose between P21 and P56, sEPSC interevent intervals are similar at both ages. While counterintuitive, this functional outcome is in line with what has been reported previously in dorsal striatum^45^ and suggests that inputs from thalamus and cortex may be weighted differently over time or that changes in release probability are actively compensating as new synapses are formed during this epoch^55^. Spontaneous EPSC amplitudes were also similar at both ages, but selective stimulation of PFC axon terminals showed that cortically evoked responses increased in strength between P21 and P56. The increase is likely due to increases in AMPA and kainate glutamate receptor subtype-mediated responses observed at corticostriatal synapses through the fourth postnatal week^45^, but also suggests there may be differences in the maturational time course for particular inputs^45^. As expected, over the same postnatal period, we observed a significant decrease in intrinsic excitability, with rheobase increasing and input resistance declining^45,46^. Rbp4-Cre mediated, presynaptic deletion of Cdh8 had no impact on the generation of corticostriatal synaptic appositions but did produce a lasting reduction in sEPSC frequency.

Our data also contribute to evidence showing that adhesion between classic and type II cadherins promotes appropriate pre- to postsynaptic matching. PFC axons lacking Cdh8 shift their target fields ventrally in striatum in a manner similar to data showing that retinal bipolar cells lacking Cdh8 extend their axons beyond their normal target layer^8^. In most cases, type II cadherins have been shown to generate a high level of synapse specificity by acting combinatorially^8,56–60^ and we would predict an even greater impact on PFC axon targeting with simultaneous manipulation of several cadherins. Likely candidates for Cdh8 collaboration in dorsal striatum are Cdh11, which can bind Cdh8 heterophilically and is expressed in a partially overlapping pattern in PFC and striatum^9,11,61,62^ and N-cadherin^59^. These actions would be expected to occur in the context of complementary actions driven by other molecular interactions, such as those mediated by Pcdh17, which has an expression pattern that overlaps that of Cdh8, and which may be important for cortical neuron development^63,64^, and Sema3E, PlexinD1 and Ten-m3, which support thalamostriatal synapse targeting^65,66^.

In contrast to shifting the position of corticostriatal axonal arbors, we observed no change in the density of corticostriatal (VGLUT1-expressing) synapse appositions or in the density of SPN dendritic protrusions suggesting that mistargeted axons form synapses. That loss of Cdh8 does not impede synaptogenesis is consistent with data from Cdh8 knockout mice where peripheral sensory neurons that normally express Cdh8 can form synapses, but display reduced cold sensitivity^67^, and mistargeted retinal bipolar cell axons form synapses that fail to encode OFF visual responses^8^. These data also support the idea that the impact of Cdh8 loss may be magnified by both losing appropriate connections and gaining inappropriate ones, as evidenced here by the greatly impaired performance of Cdh8 cKO mice on an instrumental learning task.

Finally, we also generated and applied a novel strategy, Hotspot Analysis, to identify and track patterns in a brain region containing few anatomical landmarks (like layers) and one that is also expanding in size. Key advantages of Hotspot are its simplicity and z-score based readouts that can be used for conducting screens aimed at detecting altered connectivity that could be further pursued in future studies using high resolution approaches that require very large datasets and computational power.

## Author contributions

RM, DLB and GWH conceived the study. RM, YZ, DLB, GWH designed the experiments. Methodology and implementation: stereology (RM); Hotspot development (YZ); Hotspot analysis (RM, YZ, DLB), synapse analysis (RM, DLB); floxed mouse (LGF, RM), behavior (AH); electrophysiology (CAG); neuron reconstruction (RM, CAG, KT); anatomy and imaging pipeline (RM, YZ, PDV, ARM, NT); analysis (RM, YZ, DLB, GWH, MGB). RM, DLB, GWH, YZ, MGB, AH contributed to interpretation of results. Writing-original draft: RM and DLB; Writing-review and editing: RM, YZ, DLB, GWH, MGB; LGF, KT, AM, NT.

## Acknowledgments

We thank Frances Williams, Romario Thomas, and Mofida Abdelmageed for technical support. We would like to thank Dr. Sijie Hao for help in developing analysis strategies, and to acknowledge the ISMMS Microscopy and Advanced Bioimaging Core for access to instruments and expertise and the ISMMS Mouse Genetics and Gene Targeting Core and Dr. Kevin Kelley for reviving the Rpb4Cre line and for generating and cryopreserving the *Cdh8* conditional mutant mouse.

## Inclusion and diversity

One or more of the authors of this paper self-identifies as a group that is underrepresented in science. One or more of the authors of this paper self-identifies as a member of the LGBTQIA+ community. One or more of the authors of this paper self-identifies as a gender minority in their field of research. One or more of the authors of this paper received support from a program designed to increase minority representation in science.

## Methods

### RESOURCE AVAILABILITY

#### Lead contact

Further information and requests for resources and reagents should be directed to and will be fulfilled by the lead contact, DLB.

#### Materials availability

Cdh8^fl/fl^ line generated in this study has been deposited at Jackson Labs [JAX: 037592].

#### Data and code availability

All original code has been deposited at GitHub and is publicly available.

### EXPERIMENTAL MODEL AND SUBJECT DETAILS

Mice of both sexes from the following lines were used to generate data: C57BL/6J (WT) mice (IMSR_JAX:000664); Cadherin-8 floxed mice (Cdh8^fl/fl/^; see below); RBP4-Cre mice (MMRRC_037128; GENSAT); Ai9 Cre reporter mice (IMSR_JAX:007909); and NEX-Cre mice (generous gift from Klaus-Armin Nave, Max Planck, Göttingen ^26^. All animals were kept with dams until weaning age (P21) and then housed in single sex groups of 3-5 animals per cage. The care and treatment of all animals were in strict accordance with guidelines of the Institutional Animal Care and Use Committee of the ISMMS and those of the National Institutes of Health.

The Cdh8^fl/fl/^ mice were generated through embryonic stem cell germline transmission using a construct obtained from UC Davis, KOMP Repository: Cdh8^tm2a^KOMP^Wtsi 69^. ES cells were injected into C57BL/6J blastocysts (Jackson Laboratories) at the ISMMS Mouse Genetics Core. The *Cdh8* construct contains a lacZ cassette adjacent to a Neo cassette that are flanked by FRT sites and exon 3 of *Cdh8*, flanked by LoxP sites. Insertion and germline transmission in progeny was confirmed by long-range PCR. Site of insertion was also consistent with the expression pattern of LacZ staining, which matched published studies of Cdh8 mRNA distribution ^10,11^. The Neo (and LacZ) cassette was then removed by crossing the Cdh8^fl/fl^ mice with a Flp recombinase line (IMSR_JAX:012930) and its deletion was confirmed by PCR. For this and all crosses, genotype was assessed with tail DNA extracted using RED Extract-N-Amp PCR Ready Mix (Sigma-Aldrich) followed by a polymerase chain reaction (PCR) using the relevant primers (Table 1). Amplicons were then visualized in a 1.5% agarose gel with ethidium bromide. Mice (Cdh8^tm2c^ have been deposited at Jackson labs (JAX: 037592).

## METHOD DETAILS

### Stereotaxic Surgeries

Mice were anesthetized by hypothermia (P0-P1 pups) or with continuous delivery of isoflurane (P14 or older) and mounted in a stereotaxic frame (David Kopf Instruments, Tujunga, CA). Mice were injected with AAV1-hSyn-TurboRFP or -mGFP (Addgene or UPenn Vector Core; 105552; gift from James M Wilson); 50465; gift from Bryan Roth) at P0, P14 or P49 (7 days prior to sacrifice). For Rbp4-Cre mice, pAAV1 hSyn FLEx mGFP-2A-Synaptophysin-mRuby (Stanford Vector Core; gift from Liqun Luo^70^) was introduced at P7 or P42 (14-days prior to sacrifice). For electrophysiological studies, PFC-neurons were transduced with pAAV1-CaMKIIa-hCHR2(H134R)- EYFP (Addgene 26969; gift from Karl Deisseroth^71^) 14-days prior to recording. Virus was delivered unilaterally using a Drummond Nanoject III at a speed of 2nl/sec, into the PFC using the following coordinates from bregma: P0: A/P= 0.3, M/L= ± 0.1, D/V= -0.9; P14: A/P= 1.4, M/L= ± 0.3, D/V = - 1.6; P49: A/P=1.9, M/L= ± 0.3, D/V = -2.2 and total volume according brain size and titer (adapted from^72^ or S1: A/P = -1.5, M/L = ±3.0, D/V = -0.7.

### Tissue Preparation

Animals were perfused transcardially with saline (P7) or 1% paraformaldehyde (P21, P56) to clear vasculature, followed by ice chilled 4% paraformaldehyde in phosphate-buffered saline (PBS), pH7.2 for 10 min. Brains were post-fixed for 10-14 hrs at 4°C, cryoprotected in 7% sucrose/PBS and sectioned coronally using a vibratome (VT1000S, Leica Biosystems) at a setting of 50 µm through the entire extent of the striatum. All sections were collected and kept in order.

### Stereology

For systematic random sampling, every 5^th^ section was sampled beginning at the caudal end of the olfactory bulb to the anterior hippocampus. Tissue was washed in 1x PBS, mounted in VectaShield with DAPI (Vector Labs, H1200) and visualized on a Zeiss AxioImager widefield microscope using a 100X, 1.4NA objective lens. Using StereoInvestigator (MBF Bioscience), axon length was estimated by counting axons crossing a virtual spaceball hemisphere (radius: 3 µm) within a guard zone height of 10 µm and a grid size of 200 µm × 300 µm (3-5 axons per grid) mapped in a systematically random fashion across striatum. The PFC was defined as the anterior cingulate (ACC), prelimbic (PL) and infralimbic (IL) cortex (Fig. 1B). Regions of interest (PFC, injection site, and striatum) were delineated using the Nissl (DAPI) staining pattern and the Allen Brain Mouse Atlas as a reference. Volumes were measured using the Cavalieri Estimator probe (MBF) and Cavalieri_V_estimator (Image J) (point spacing 140 µm) on stitched tiled images acquired on a Zeiss LSM780 confocal microscope using a 20X objective lens.

### Hotspot Analysis

Sections were sampled as above and image tiles were captured and stitched using a Zeiss LSM 780 or 980 confocal microscope and a 20x objective. Eight-bit images were exported as 8-bit tifs to generate ROIs using FIJI. For Hotspot Analysis, we utilized the Getis Ord Gi* Statistic^7^. This statistical method calculates the density of pixel values within a neighborhood, Gi, and the expected density of pixel values under the null hypothesis, E(Gi). It then uses those two values along with the variance of the dataset to generate a Z-score, which can be used to discern the likelihood that the observed level of pixel clustering within a neighborhood has occurred by chance. The Gi statistic is explained below:

The expected value of a given pixel under the null hypothesis of spatial randomness is the global mean pixel value:

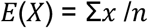

where:

*x* are pixel values

*n* are the number of pixels in an image

Then, the expected Gi statistic within any given neighborhood is:

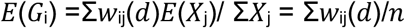

where:

*w*_ij_(*d*) are all the pixels *w* within distance *j* of pixel *i*

Σ*X*j is the sum of all pixel values *j* within distance *d*

The actual density of pixel values within a neighborhood is calculated as: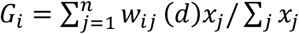

where:

*x*_j_ represents all the pixels in the neighborhood

The Z-score comparing the expected and actual Gi statistics is calculated as:

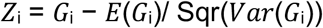

where:

*Var*(*G*_i_) is the variance of the *G*_i_ statistic

By delineating an ROI, the dorsal striatum, and removing by thresholding the saturated pixels that comprise the bundles of descending axons in the internal capsule, maps were created by using the Gi statistic associated with a 20×20 pixel-sized neighborhood (local mean; size based on results from pilot studies). The output of this Python script provided a visual representation of axon clustering through the striatum and the summary statistics representing the z-scores for each pixel neighborhood. The custom scripts used for Hotspot Analysis can be found at: github.com/deannabenson/HotspotAnalysis.

Basic geometry was used to compare distances between XY coordinates at max Gi. In Excel this translates to: {=MIN(SQRT(((A1-D$1:D$z)^2)+((B1-E$1:E$z)^2)))}, where X values are in columns A and D; Y values in columns B and E.

### Immunohistochemistry and Synapse Analysis

Tissue sections collected as for stereological studies were permeabilized using 0.25% Triton-X-100 (Sigma) for 5-7 min (RT), blocked with 5% normal donkey serum (Jackson ImmunoResearch) for 1h, RT, all in PBS and incubated sequentially in primary and secondary antibodies: first in chicken anti-Homer 1 (1:500, Synaptic Systems, Cat#160 006)/2% serum/PBS, visualized with anti-chicken Alexa-488 (1:200, Jackson ImmunoResearch); and then by guinea pig anti-vesicular glutamate transporter 1 antibody (V-GLUT1, 1:1000, Millipore-Sigma, Cat#AB5905)/2% serum, visualized with anti-guinea pig Alexa-647 and then coverslipped in Vectashield, and sealed with nail polish. Single, optical confocal images were acquired using a 100x, 1.4NA objective set at a zoom factor of 2 using an upright, fixed-stage Zeiss LSM780 (resolution of 24 pixels per micron). Images were captured across three regions in striatum (dorsal, lateral, ventral; Fig. 5B); five images per section, four sections per animal, and 3-5 animals per group. Images were imported into FIJI, and puncta were segmented by thresholding and used to generate masks. Puncta size and density (number per area) were assessed using “analyze particles” function set to exclude puncta below 0.07µm^2^, a value determined in pilot studies and maintained throughout. Appositions were assessed by multiplying VGLUT1 and Homer image masks using the “image calculator” function, setting all unmatched pixels to zero.

### Western Blot

Whole-cell lysates were generated from cortex or striatum (10mM HEPES, 2mM EDTA, 2mM EGTA, 0.5 mM DTT, phosphatase and protease inhibitor cocktails (Sigma) via 3 × 5s pulses using a Kontes pestle gun. Samples were centrifuged 1500g, 10mins, 4°C and protein concentrations were assessed using a Bradford assay (BioRad). Proteins (25µg) per lane were separated using 8% SDS-PAGE, transferred to Immunoblot PVDF membranes (Millipore), blocked with LI-COR Odyssey Blocking Buffer with 0.1% Tween 20 (Thermo), incubated with primary antibodies directed against Cdh8 (sc6461, Santa Cruz), N-cadherin (BD Transduction Labs; 610921), ß-catenin (Millipore; 06734), ß-tubulin (Abcam; Ab125267). Membranes were washed and incubated with DyLight 800 or DyLight 680 conjugated secondaries (Pierce) and then imaged using a LI-COR Odyssey CLX imager (LI-COR Biosciences).

### Electrophysiology

Whole-cell patch-clamp recordings from Cdh8^fl/fl^ or Cdh8 cKO spiny projection neurons (SPNs) in the dorsal medial striatum were conducted in acute coronal slices (350 µm-thick) taken after deep anesthesia with isoflurane and rapid decapitation at P21 or P56. After 1h incubation at 37°C, slices were placed in an immersion chamber containing gabazine (GBZ; 10 µM; Sigma) at 31°C, which was used for all electrophysiological experiments. Recordings were made with a Multiclamp 700B amplifier (Molecular Devices, Sunnyvale, CA). Analog signals were low-pass filtered, digitized, and analyzed with pClamp10 software (Molecular Devices, Sunnyvale, CA). SPNs were visualized with an infrared camera (IR-1000; DAGE MTI) mounted on an upright epifluorescence microscope (BX50WI; Olympus). Whole-cell recordings were acquired using glass micropipettes with a resistance of 2–4 MΩ and filled with an intracellular solution containing 124mM K-gluconate, 10mM HEPES, 10mM phosphocreatine di(Tris), 0.2mM EGTA, 4mM Mg2ATP, 0.3mM Na2GTP, and (0.3%) biocytin. SPNs were recorded for 3 min following a 5 min equilibration period. After recording sEPSC activity in voltage-clamp mode, SPNs were identified electrophysiologically in current-clamp mode by observing characteristic patterns of action potentials induced by a series of depolarizing current pulses (10-s interpulse intervals, 500-ms-long pulses, 20-pA current intervals) (Fino et al. 2007, Matikainen-Ankney et al. 2016; Guevara et al. 2020). Recordings of SPN firing characteristics were used for analysis in Clampfit for properties of intrinsic excitability. Input-output (I/O) experiments were conducted on acute slices collected from P21 and P56 WT mice that had been injected with an EYFP-tagged channelrhodopsin-2 virus in the PFC 14-days prior to recordings^73^. PFC-terminals surrounding SPNs in DMS were stimulated utilizing a transistor–transistor logic (TTL)-pulsed microscope objective-coupled LEDs (Prizmatix). Input-output curves were generated by increasing light intensity in a step- wise manner (0.2mW/mm–2 per step), from the minimal stimulation level 1 at resting membrane potential in order to evoke glutamatergic mediated responses. The elicited power of light stimuli was measured using an optical power meter (Thorlabs). For each stimulus, optically-evoked EPSC (oEPSC) amplitudes were normalized to the evoked amplitude response of the first (minimal) stimulation intensity for every individual cell. This was to account for the impact of variations in viral expression levels as described ^74^.

### Morphometric analysis of biocytin-filled neurons

General procedures and spine morphometric analyses followed our general previously published protocols^75,76^. All recorded SPNs were filled with biocytin (Tocris Bioscience) contained in the patch pipette. After recording, slices were immersed in 4% paraformaldehyde overnight at 4°C, permeabilized with 0.1% Triton X-100 for 2h at RT, and labeled with streptavidin-conjugated Alexa Fluor 594 (1:300; Jackson ImmunoResearch) at 4°C for 48h. SPNs were then imaged on an upright, fixed stage, confocal LSM 780 (Zeiss) using a 63x 1.4N.A.objective, and Nyquist sampling to visualize and render dendritic protrusions (0.067 × 0.067 × 0.474 µm per voxel). Z-stacks were deconvolved using AutoDeblur/AutoVisualize X (MediaCybernetics). Dendrite protrusion density and morphology were assessed using NeuronStudio^77^.

### Behavior

Instrumental learning was carried out in Bussey Saksida touchscreen-based operant chambers (Lafayette Instruments) using a protocol established previously (Hussein et al, 2022) in which mice were trained to nosepoke a single, lit response window to obtain a reward (strawberry milk). The presence of a reward was cued by the presentation of a tone. Two 60-minute training sessions served to habituate mice to the apparatus, and this was followed by a 2h continuous reinforcement schedule (CRF) in which each nose poke was rewarded with a reward. Mice then progressed to the testing phase which each 60m session was administered over four consecutive days. The first session was CRF, and the remaining three were random interval (RI) schedules in which the probability of the reward remained constant over a 15s (day 2) 30s (day 3) or 30s (day 4) interval. The random interval timer was initiated by the first nose poke. Schedules were controlled and data were collected using ABET II Touch software (Lafayette) and then exported to Excel.

## QUANTIFICATION AND STATISTICAL ANALYSES

Both sexes were used, and data were analyzed for sex differences. Because no sex-dependent differences were found, data from both males and females were pooled with the exception of behavior, where only male mice were used. Group sizes were 3-5 mice per genotype for anatomical and synaptic studies (5 images per region; 3-5 sections per animal) and ~15 cells per condition, from 8-15 brain slices, from 5-6 mice per genotype/sex. Where relevant, genotype identity was coded and analyzed blind. Data were expressed as mean + SEM. Repeated measures ANOVA with post hoc Bonferroni corrections for multiple comparisons and 2-way ANOVA or mixed-effects model analysis were used to delineate impact of the manipulation vs. genotype or age. Unpaired Student’s t-tests and Mann-Whitney U (Wilcoxon’s rank sum) test were used for comparison between conditions and cell types for electrophysiology experiments. For analysis of dendritic protrusions, groups were compared using 1-way ANOVA. Spine-head diameter and spine length cumulative frequency distributions were compared using two-sample Kolmogorov–Smirnov tests. For all experiments, a minimum criterion of p< 0.05 was used for biological significance. Additional specifics are given in figure legends.

## REFERENCES

1. Balleine, B.W., Delgado, M.R., and Hikosaka, O. (2007). The role of the dorsal striatum in reward and decision-making. J Neurosci 27, 8161–8165. 10.1523/JNEUROSCI.1554-07.2007.

2. Kennerley, S.W., Walton, M.E., Behrens, T.E.J., Buckley, M.J., and Rushworth, M.F.S. (2006). Optimal decision making and the anterior cingulate cortex. Nat Neurosci 9, 940–947. 10.1038/nn1724.

3. Klune, C.B., Jin, B., and DeNardo, L.A. (2021). Linking mPFC circuit maturation to the developmental regulation of emotional memory and cognitive flexibility. eLife 10, e64567. 10.7554/eLife.64567.

4. Li, W., and Pozzo-Miller, L. (2020). Dysfunction of the corticostriatal pathway in autism spectrum disorders. J Neurosci Res 98, 2130–2147. 10.1002/jnr.24560.

5. Shepherd, G.M.G. (2013). Corticostriatal connectivity and its role in disease. Nat Rev Neurosci 14, 278–291. 10.1038/nrn3469.

6. Bayer, S.A., and Altman, J. (1987). Directions in neurogenetic gradients and patterns of anatomical connections in the telencephalon. Prog Neurobiol 29, 57–106. 10.1016/0301-0082(87)90015-3.

7. Getis, A., and Ord, J.K. (1992). The Analysis of Spatial Association by Use of Distance Statistics. Geographical Analysis 24, 189–206. 10.1111/j.1538-4632.1992.tb00261.x.

8. Duan, X., Krishnaswamy, A., De la Huerta, I., and Sanes, J.R. (2014). Type II cadherins guide assembly of a direction-selective retinal circuit. Cell 158, 793–807. 10.1016/j.cell.2014.06.047.

9. Frei, J.A., Niescier, R.F., Bridi, M.S., Durens, M., Nestor, J.E., Kilander, M.B.C., Yuan, X., Dykxhoorn, D.M., Nestor, M.W., Huang, S., et al. (2021). Regulation of Neural Circuit Development by Cadherin-11 Provides Implications for Autism. eNeuro 8, ENEURO.0066-21.2021. 10.1523/ENEURO.0066-21.2021.

10. Friedman, L.G., Riemslagh, F.W., Sullivan, J.M., Mesias, R., Williams, F.M., Huntley, G.W., and Benson, D.L. (2014). Cadherin-8 expression, synaptic localization, and molecular control of neuronal form in prefrontal corticostriatal circuits. The Journal of comparative neurology. 10.1002/cne.23666.

11. Suzuki, S.C., Inoue, T., Kimura, Y., Tanaka, T., and Takeichi, M. (1997). Neuronal circuits are subdivided by differential expression of type-II classic cadherins in postnatal mouse brains. Mol. Cell Neurosci. 9, 433–447.

12. Marchand, R., and Lajoie, L. (1986). Histogenesis of the striopallidal system in the rat. Neurogenesis of its neurons. Neuroscience 17, 573–590. 10.1016/0306-4522(86)90031-x.

13. Deacon, T.W., Pakzaban, P., and Isacson, O. (1994). The lateral ganglionic eminence is the origin of cells committed to striatal phenotypes: neural transplantation and developmental evidence. Brain Res 668, 211–219. 10.1016/0006-8993(94)90526-6.

14. van der Kooy, D., and Fishell, G. (1987). Neuronal birthdate underlies the development of striatal compartments. Brain Res 401, 155–161. 10.1016/0006-8993(87)91176-0.

15. Sohur, U.S., Padmanabhan, H.K., Kotchetkov, I.S., Menezes, J.R., and Macklis, J.D. (2014). Anatomic and molecular development of corticostriatal projection neurons in mice. Cereb Cortex 24, 293–303. 10.1093/cercor/bhs342.

16. Balleine, B.W., and O’Doherty, J.P. (2010). Human and Rodent Homologies in Action Control: Corticostriatal Determinants of Goal-Directed and Habitual Action. Neuropsychopharmacology 35, 48–69. 10.1038/npp.2009.131.

17. Rushworth, M.F.S., Walton, M.E., Kennerley, S.W., and Bannerman, D.M. (2004). Action sets and decisions in the medial frontal cortex. Trends in Cognitive Sciences 8, 410–417. 10.1016/j.tics.2004.07.009.

18. Calhoun, M.E., Mao, Y., Roberts, J.A., and Rapp, P.R. (2004). Reduction in hippocampal cholinergic innervation is unrelated to recognition memory impairment in aged rhesus monkeys. The Journal of comparative neurology 475, 238–246. 10.1002/cne.20181.

19. Van Eden, C.G., and Uylings, H.B. (1985). Postnatal volumetric development of the prefrontal cortex in the rat. J Comp Neurol 241, 268–274. 10.1002/cne.902410303.

20. Hunnicutt, B.J., Jongbloets, B.C., Birdsong, W.T., Gertz, K.J., Zhong, H., and Mao, T. (2016). A comprehensive excitatory input map of the striatum reveals novel functional organization. Elife 5. 10.7554/eLife.19103.

21. Hintiryan, H., Foster, N.N., Bowman, I., Bay, M., Song, M.Y., Gou, L., Yamashita, S., Bienkowski, M.S., Zingg, B., Zhu, M., et al. (2016). The mouse cortico-striatal projectome. Nat Neurosci 19, 1100–1114. 10.1038/nn.4332.

22. Groenewegen, H.J., Vermeulen-Van der Zee, E., te Kortschot, A., and Witter, M.P. (1987). Organization of the projections from the subiculum to the ventral striatum in the rat. A study using anterograde transport of Phaseolus vulgaris leucoagglutinin. Neuroscience 23, 103–120. 10.1016/0306-4522(87)90275-2.

23. Kelley, A.E., Domesick, V.B., and Nauta, W.J. (1982). The amygdalostriatal projection in the rat--an anatomical study by anterograde and retrograde tracing methods. Neuroscience 7, 615–630. 10.1016/0306-4522(82)90067-7.

24. Fentress, J.C., Stanfield, B.B., and Cowan, W.M. (1981). Observation on the development of the striatum in mice and rats. Anat Embryol (Berl) 163, 275–298. 10.1007/BF00315705.

25. Sharpe, N.A., and Tepper, J.M. (1998). Postnatal development of excitatory synaptic input to the rat neostriatum: an electron microscopic study. Neuroscience 84, 1163–1175.

26. Goebbels, S., Bormuth, I., Bode, U., Hermanson, O., Schwab, M.H., and Nave, K.A. (2006). Genetic targeting of principal neurons in neocortex and hippocampus of NEX-Cre mice. Genesis 44, 611–621. 10.1002/dvg.20256.

27. Memi, F., Killen, A.C., Barber, M., Parnavelas, J.G., and Andrews, W.D. (2019). Cadherin 8 regulates proliferation of cortical interneuron progenitors. Brain Struct Funct 224, 277–292. 10.1007/s00429-018-1772-4.

28. Harris, J.A., Mihalas, S., Hirokawa, K.E., Whitesell, J.D., Choi, H., Bernard, A., Bohn, P., Caldejon, S., Casal, L., Cho, A., et al. (2019). Hierarchical organization of cortical and thalamic connectivity. Nature 575, 195–202. 10.1038/s41586-019-1716-z.

29. Gerfen, C.R., Paletzki, R., and Heintz, N. (2013). GENSAT BAC cre-recombinase driver lines to study the functional organization of cerebral cortical and basal ganglia circuits. Neuron 80, 1368– 1383. 10.1016/j.neuron.2013.10.016.

30. Kozorovitskiy, Y., Saunders, A., Johnson, C.A., Lowell, B.B., and Sabatini, B.L. (2012). Recurrent network activity drives striatal synaptogenesis. Nature 485, 646–650. 10.1038/nature11052.

31. Leone, D.P., Heavner, W.E., Ferenczi, E.A., Dobreva, G., Huguenard, J.R., Grosschedl, R., and McConnell, S.K. (2015). Satb2 Regulates the Differentiation of Both Callosal and Subcerebral Projection Neurons in the Developing Cerebral Cortex. Cereb Cortex 25, 3406–3419. 10.1093/cercor/bhu156.

32. Lein, E.S., Hawrylycz, M.J., Ao, N., Ayres, M., Bensinger, A., Bernard, A., Boe, A.F., Boguski, M.S., Brockway, K.S., Byrnes, E.J., et al. (2007). Genome-wide atlas of gene expression in the adult mouse brain. Nature 445, 168–176. 10.1038/nature05453.

33. Bozdagi, O., Wang, X.B., Nikitczuk, J.S., Anderson, T.R., Bloss, E.B., Radice, G.L., Zhou, Q., Benson, D.L., and Huntley, G.W. (2010). Persistence of coordinated long-term potentiation and dendritic spine enlargement at mature hippocampal CA1 synapses requires N-cadherin. J Neurosci 30, 9984–9989. 10.1523/JNEUROSCI.1223-10.2010.

34. Huntley, G.W., Elste, A.M., Patil, S.B., Bozdagi, O., Benson, D.L., and Steward, O. (2012). Synaptic loss and retention of different classic cadherins with LTP-associated synaptic structural remodeling in vivo. Hippocampus 22, 17–28. 10.1002/hipo.20859.

35. Hussein, A., Tielemans, A., Baxter, M.G., Benson, D.L., and Huntley, G.W. (2022). Cognitive deficits and altered cholinergic innervation in young adult male mice carrying a Parkinson’s disease Lrrk2G2019S knockin mutation. Exp Neurol 355, 114145. 10.1016/j.expneurol.2022.114145.

36. Shan, Q., Ge, M., Christie, M.J., and Balleine, B.W. (2014). The acquisition of goal-directed actions generates opposing plasticity in direct and indirect pathways in dorsomedial striatum. J Neurosci 34, 9196–9201. 10.1523/JNEUROSCI.0313-14.2014.

37. Shiflett, M.W., Brown, R.A., and Balleine, B.W. (2010). Acquisition and performance of goal-directed instrumental actions depends on ERK signaling in distinct regions of dorsal striatum in rats. J Neurosci 30, 2951–2959. 10.1523/JNEUROSCI.1778-09.2010.

38. Agmon, A., Yang, L.T., Jones, E.G., and O’Dowd, D.K. (1995). Topological precision in the thalamic projection to neonatal mouse barrel cortex. J Neurosci 15, 549–561.

39. Simpson, H.D., Kita, E.M., Scott, E.K., and Goodhill, G.J. (2013). A quantitative analysis of branching, growth cone turning, and directed growth in zebrafish retinotectal axon guidance. J Comp Neurol 521, 1409–1429. 10.1002/cne.23248.

40. Riccomagno, M.M., and Kolodkin, A.L. (2015). Sculpting neural circuits by axon and dendrite pruning. Annu Rev Cell Dev Biol 31, 779–805. 10.1146/annurev-cellbio-100913-013038.

41. Tran, H. (2013). The role of Ten-m3 in the development of the mouse thalamostriatal pathway.

42. Mandelbaum, G., Taranda, J., Haynes, T.M., Hochbaum, D.R., Huang, K.W., Hyun, M., Umadevi Venkataraju, K., Straub, C., Wang, W., Robertson, K., et al. (2019). Distinct Cortical-Thalamic-Striatal Circuits through the Parafascicular Nucleus. Neuron 102, 636-652.e7. 10.1016/j.neuron.2019.02.035.

43. Uryu, K., Butler, A.K., and Chesselet, M.F. (1999). Synaptogenesis and ultrastructural localization of the polysialylated neural cell adhesion molecule in the developing striatum. J Comp Neurol 405, 216–232. 10.1002/(sici)1096-9861(19990308)405:2<216::aid-cne6>3.0.co;2-6.

44. Hattori, T., and McGeer, P.L. (1973). Synaptogenesis in the corpus striatum of infant rat. Exp Neurol 38, 70–79. 10.1016/0014-4886(73)90008-3.

45. Krajeski, R.N., Macey-Dare, A., van Heusden, F., Ebrahimjee, F., and Ellender, T.J. (2019). Dynamic postnatal development of the cellular and circuit properties of striatal D1 and D2 spiny projection neurons. J Physiol 597, 5265–5293. 10.1113/JP278416.

46. Peixoto, R.T., Wang, W., Croney, D.M., Kozorovitskiy, Y., and Sabatini, B.L. (2016). Early hyperactivity and precocious maturation of corticostriatal circuits in Shank3B(-/-) mice. Nat Neurosci 19, 716–724. 10.1038/nn.4260.

47. Savage, J.C., St-Pierre, M.-K., Carrier, M., El Hajj, H., Novak, S.W., Sanchez, M.G., Cicchetti, F., and Tremblay, M.-È. (2020). Microglial physiological properties and interactions with synapses are altered at presymptomatic stages in a mouse model of Huntington’s disease pathology. J Neuroinflammation 17, 98. 10.1186/s12974-020-01782-9.

48. Tepper, J.M., Sharpe, N.A., Koos, T.Z., and Trent, F. (1998). Postnatal development of the rat neostriatum: electrophysiological, light- and electron-microscopic studies. Developmental neuroscience 20, 125–145.

49. Cline, H.T. (2001). Dendritic arbor development and synaptogenesis. Current opinion in neurobiology 11, 118–126.

50. Rakic, P., Bourgeois, J.P., Eckenhoff, M.F., Zecevic, N., and Goldman-Rakic, P.S. (1986). Concurrent overproduction of synapses in diverse regions of the primate cerebral cortex. Science 232, 232–235. 10.1126/science.3952506.

51. Wilton, D.K., Dissing-Olesen, L., and Stevens, B. (2019). Neuron-Glia Signaling in Synapse Elimination. Annu Rev Neurosci 42, 107–127. 10.1146/annurev-neuro-070918-050306.

52. Johnson, C.M., Peckler, H., Tai, L.-H., and Wilbrecht, L. (2016). Rule learning enhances structural plasticity of long-range axons in frontal cortex. Nat Commun 7, 10785. 10.1038/ncomms10785.

53. Ingham, C.A., Hood, S.H., Taggart, P., and Arbuthnott, G.W. (1998). Plasticity of Synapses in the Rat Neostriatum after Unilateral Lesion of the Nigrostriatal Dopaminergic Pathway. J Neurosci 18, 4732–4743. 10.1523/JNEUROSCI.18-12-04732.1998.

54. McNeill, T.H., Brown, S.A., Rafols, J.A., and Shoulson, I. (1988). Atrophy of medium spiny I striatal dendrites in advanced Parkinson’s disease. Brain Res 455, 148–152. 10.1016/0006-8993(88)90124-2.

55. Choi, S., and Lovinger, D.M. (1997). Decreased probability of neurotransmitter release underlies striatal long-term depression and postnatal development of corticostriatal synapses. Proc Natl Acad Sci U S A 94, 2665–2670.

56. Basu, R., Duan, X., Taylor, M.R., Martin, E.A., Muralidhar, S., Wang, Y., Gangi-Wellman, L., Das, S.C., Yamagata, M., West, P.J., et al. (2017). Heterophilic Type II Cadherins Are Required for High-Magnitude Synaptic Potentiation in the Hippocampus. Neuron 96, 160-176.e8. 10.1016/j.neuron.2017.09.009.

57. Kuwako, K., Nishimoto, Y., Kawase, S., Okano, H.J., and Okano, H. (2014). Cadherin-7 Regulates Mossy Fiber Connectivity in the Cerebellum. Cell Reports 9, 311–323. 10.1016/j.celrep.2014.08.063.

58. Osterhout, J.A., Josten, N., Yamada, J., Pan, F., Wu, S.W., Nguyen, P.L., Panagiotakos, G., Inoue, Y.U., Egusa, S.F., Volgyi, B., et al. (2011). Cadherin-6 mediates axon-target matching in a non-image-forming visual circuit. Neuron 71, 632–639. 10.1016/j.neuron.2011.07.006.

59. Vagnozzi, A.N., Moore, M.T., Lin, M., Brozost, E.M., Kc, R., Agarwal, A., Schwarz, L.A., Duan, X., Zampieri, N., Landmesser, L.T., et al. (2022). Coordinated cadherin functions sculpt respiratory motor circuit connectivity. Elife 11, e82116. 10.7554/eLife.82116.

60. Duan, X., Krishnaswamy, A., Laboulaye, M.A., Liu, J., Peng, Y.-R., Yamagata, M., Toma, K., and Sanes, J.R. (2018). Cadherin Combinations Recruit Dendrites of Distinct Retinal Neurons to a Shared Interneuronal Scaffold. Neuron 99, 1145-1154.e6. 10.1016/j.neuron.2018.08.019.

61. Brasch, J., Katsamba, P.S., Harrison, O.J., Ahlsén, G., Troyanovsky, R.B., Indra, I., Kaczynska, A., Kaeser, B., Troyanovsky, S., Honig, B., et al. (2018). Homophilic and Heterophilic Interactions of Type II Cadherins Identify Specificity Groups Underlying Cell-Adhesive Behavior. Cell Rep 23, 1840–1852. 10.1016/j.celrep.2018.04.012.

62. Patel, S.D., Ciatto, C., Chen, C.P., Bahna, F., Rajebhosale, M., Arkus, N., Schieren, I., Jessell, T.M., Honig, B., Price, S.R., et al. (2006). Type II cadherin ectodomain structures: implications for classical cadherin specificity. Cell 124, 1255–1268.

63. Chang, H., Hoshina, N., Zhang, C., Ma, Y., Cao, H., Wang, Y., Wu, D.-d, Bergen, S.E., Landén, M., Hultman, C.M., et al. (2018). The protocadherin 17 gene affects cognition, personality, amygdala structure and function, synapse development and risk of major mood disorders. Mol Psychiatry 23, 400–412. 10.1038/mp.2016.231.

64. Hoshina, N., Tanimura, A., Yamasaki, M., Inoue, T., Fukabori, R., Kuroda, T., Yokoyama, K., Tezuka, T., Sagara, H., Hirano, S., et al. (2013). Protocadherin 17 regulates presynaptic assembly in topographic corticobasal Ganglia circuits. Neuron 78, 839–854. 10.1016/j.neuron.2013.03.031.

65. Ding, J.B., Oh, W.J., Sabatini, B.L., and Gu, C. (2012). Semaphorin 3E-Plexin-D1 signaling controls pathway-specific synapse formation in the striatum. Nature neuroscience 15, 215–223. 10.1038/nn.3003.

66. Tran, H., Sawatari, A., and Leamey, C.A. (2015). The glycoprotein Ten-m3 mediates topography and patterning of thalamostriatal projections from the parafascicular nucleus in mice. Eur J Neurosci 41, 55–68. 10.1111/ejn.12767.

67. Suzuki, S.C., Furue, H., Koga, K., Jiang, N., Nohmi, M., Shimazaki, Y., Katoh-Fukui, Y., Yokoyama, M., Yoshimura, M., and Takeichi, M. (2007). Cadherin-8 is required for the first relay synapses to receive functional inputs from primary sensory afferents for cold sensation. J Neurosci 27, 3466–3476. 10.1523/JNEUROSCI.0243-07.2007.

68. Winnubst, J., Bas, E., Ferreira, T.A., Wu, Z., Economo, M.N., Edson, P., Arthur, B.J., Bruns, C., Rokicki, K., Schauder, D., et al. (2019). Reconstruction of 1,000 Projection Neurons Reveals New Cell Types and Organization of Long-Range Connectivity in the Mouse Brain. Cell 179, 268-281.e13. 10.1016/j.cell.2019.07.042.

69. Skarnes, W.C., Rosen, B., West, A.P., Koutsourakis, M., Bushell, W., Iyer, V., Mujica, A.O., Thomas, M., Harrow, J., Cox, T., et al. (2011). A conditional knockout resource for the genome-wide study of mouse gene function. Nature 474, 337–342. 10.1038/nature10163.

70. Beier, K.T., Steinberg, E.E., DeLoach, K.E., Xie, S., Miyamichi, K., Schwarz, L., Gao, X.J., Kremer, E.J., Malenka, R.C., and Luo, L. (2015). Circuit Architecture of VTA Dopamine Neurons Revealed by Systematic Input-Output Mapping. Cell 162, 622–634. 10.1016/j.cell.2015.07.015.

71. Lee, J.H., Durand, R., Gradinaru, V., Zhang, F., Goshen, I., Kim, D.-S., Fenno, L.E., Ramakrishnan, C., and Deisseroth, K. (2010). Global and local fMRI signals driven by neurons defined optogenetically by type and wiring. Nature 465, 788–792. 10.1038/nature09108.

72. Arruda-Carvalho, M., Wu, W.-C., Cummings, K.A., and Clem, R.L. (2017). Optogenetic Examination of Prefrontal-Amygdala Synaptic Development. J Neurosci 37, 2976–2985. 10.1523/JNEUROSCI.3097-16.2017.

73. Cummings, K.A., and Clem, R.L. (2020). Prefrontal somatostatin interneurons encode fear memory. Nat Neurosci 23, 61–74. 10.1038/s41593-019-0552-7.

74. Yamamuro, K., Bicks, L.K., Leventhal, M.B., Kato, D., Im, S., Flanigan, M.E., Garkun, Y., Norman, K.J., Caro, K., Sadahiro, M., et al. (2020). A prefrontal-paraventricular thalamus circuit requires juvenile social experience to regulate adult sociability in mice. Nat Neurosci 23, 1240–1252. 10.1038/s41593-020-0695-6.

75. Guevara, C.A., Matikainen-Ankney, B.A., Kezunovic, N., LeClair, K., Conway, A.P., Menard, C., Flanigan, M.E., Pfau, M., Russo, S.J., Benson, D.L., et al. (2020). LRRK2 mutation alters behavioral, synaptic, and nonsynaptic adaptations to acute social stress. J Neurophysiol 123, 2382–2389. 10.1152/jn.00137.2020.

76. Matikainen-Ankney, B.A., Kezunovic, N., Mesias, R.E., Tian, Y., Williams, F.M., Huntley, G.W., and Benson, D.L. (2016). Altered Development of Synapse Structure and Function in Striatum Caused by Parkinson’s Disease-Linked LRRK2-G2019S Mutation. J Neurosci 36, 7128–7141. 10.1523/JNEUROSCI.3314-15.2016.

77. Rodriguez, A., Ehlenberger, D.B., Hof, P.R., and Wearne, S.L. (2006). Rayburst sampling, an algorithm for automated three-dimensional shape analysis from laser scanning microscopy images. Nat Protoc 1, 2152–2161. 10.1038/nprot.2006.313.

