## Supplemental Information for "Development of prefrontal corticostriatal connectivity in mice"

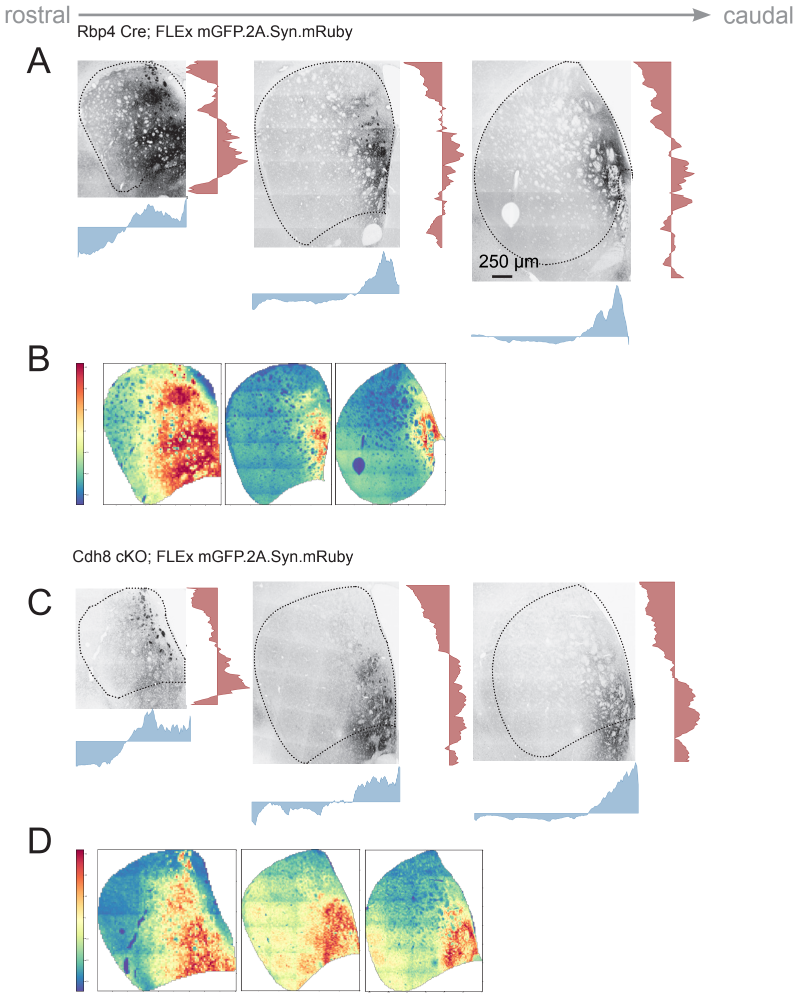


**Supplemental Figure 1**. *Presynaptic Cdh8-cKO shifts striatal territories ventrally.* Additional examples of rostral to caudal images of confocal tile scans and Hotspot Analysis through dorsal striatum of P56 Rbp4Cre control mice (A, B) and Cdh8 cKO mice (C, D) showing the distribution of axon terminals following injections of Cre-dependent Syn-mRuby. A, and C show tiled confocal images (inverted and shown in black and white so that mRuby appears black) with the dorsal striatum ROI (outline) and corresponding mountain plots. B and D show Hotspot heat maps and corresponding color scale. Magnification of confocal images shown in A.

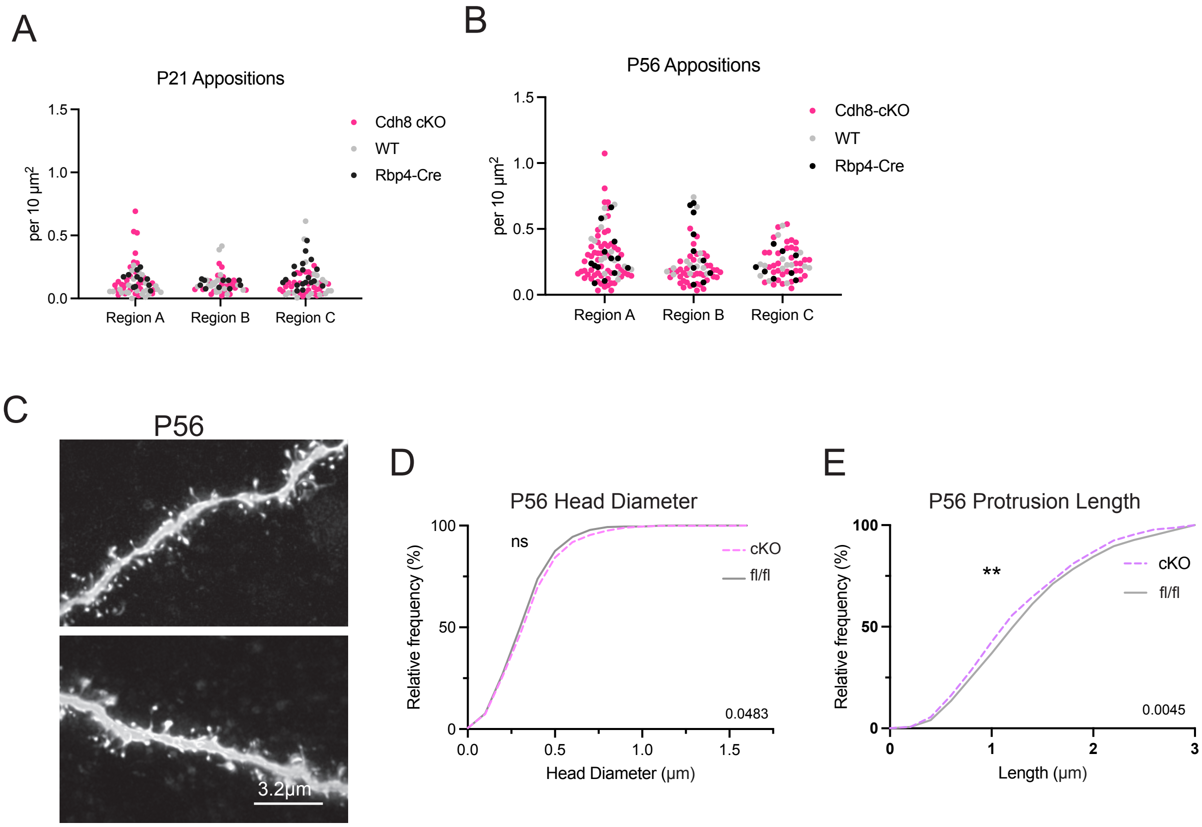


**Supplemental Figure 2.** *Cdh8 cKO shows no impact on synapse density.* Scatterplots in A and B show apposition density (vGLUT1 X Homer1) in each of the striatal regions A (medial), B (lateral) and C (ventral) defined in Figure 4. Mixed effects analyses showed no differences between genotypes within regions. Confocal images (maximum intensity projections, deconvolved) of dendritic segments from filled medial striatal projection neurons at P56 in fl/fl control (upper) and Cdh8 cKO (lower panel) mice (C). Cumulative distribution plots (D, E) showed no differences in protrusion head diameter (D), but a slight decrease in protrusion length (E) (Kolmogorov-Smirnov test; n=6-8 cells per group/ 3 dendrites per neuron/4 animals per group; **p=0.0045).

**Table 1.** *List of primers used for genotyping*

| Mouse Line | Primer | Sequence |
|---|---|---|
| | CSD-lacF (LacZ Cassette) | GCTACCATTACCAGTTGGTCTGGTGTC |
| | CSD-neoF (Neo Cassette) | GGGATCTCATGCTGGAGTTCTTCG |
| Cdh8$^{FL/FL}$ | CSD-LoxF (LoxP sites) | GAGATGGCGCAACGCAATTAATG |
| | CSD-Cdh8-R | AGCCCACCATAAAGTCATCCCATCC |
| | CSD-Cdh8-tTr | ACCAGCCTCTATAAAGTACTCAAGTTGG |
| | CSD-Cdh8-F | GCACATACCTTCACATCAAGGCTGC |
| | RBP4(31125)F | GGGCGGCCTCGGTCCTC |
| RBP4-Cre | GS Cre R2 | CCCCAGAAATGCCAGATTACGTAT |
| | Cre Up 2 (Universal Cre) | GATCTCCGGTATTGAAACTCCAGC |
| | Cre Dn 2 (Universal Cre) | GCTAAACATGCTTCATCGTCGG |
| B6 | oIMR8052 | GCG AAG AGT TTG TCC TCA ACC |
| ROSA26Flpo | oIMR8545 | AAA GTC GCT CTG AGT TGT TAT |
| (Flp mice) | oIMR8546 | GGA GCG GGA GAA ATG GAT ATG |
